# From discovery to translation: Endogenous substrates of OAT1 and OAT3 as clinical biomarkers for renal secretory function

**DOI:** 10.1101/2025.02.05.636675

**Authors:** Aarzoo Thakur, Dilip K. Singh, Katherine D. Hart, Emese Kis, Zsuzsanna Gáborik, Travis T. Denton, John D. Clarke, Mary F. Paine, Bhagwat Prasad

## Abstract

The recent ICH M12 guidance on Drug Interaction Studies encourages the use of alternate approaches for predicting drug-drug interaction (DDI) potential of new chemical entities. One approach involves biomarkers, which are endogenous substrates of drug metabolizing enzymes and transporters (DMET) and can be used to assess the inhibitory potential of new chemical entities during Phase 1 clinical studies. Thus, biomarkers could potentially eliminate the need for dedicated DDI studies with exogenous probe substrates. Metabolomics, in conjunction with *in vitro* and/or *in vivo* preclinical models or clinical studies, can be used for biomarker discovery. We developed and applied a novel metabolomics-based DMET biomarker discovery (MDBD) approach to identify and qualify biomarkers of renal organic anion transporter 1 (OAT1) and OAT3. Untargeted metabolomics of pooled plasma and urine samples from a pharmacokinetic DDI study using the OAT1/3 inhibitor, probenecid, yielded 153 features identified as putative OAT1/3 biomarkers. Subsequently, *in vitro* transporter uptake assays using processed urine samples confirmed 57 of these features as OAT1 and/or OAT3 substrates. Finally, 23 features were clinically validated as OAT1/3 biomarkers through a detailed pharmacokinetic analysis (0-24 h) of plasma and urine samples. These biomarkers, either alone or as part of a panel, can predict OAT1/3-mediated DDIs and interindividual variability in the renal secretory clearance of organic anions across different populations, thereby enabling translational utility in clinical settings. The novel MDBD approach can be extended to discover biomarkers of other transporters and enzymes.

**SUMMARY:** Using clinical and mechanistic *in vitro* approaches, 23 endogenous substrates of OAT1/3 were identified as potential clinical biomarkers of renal secretary elimination of organic anions.

## INTRODUCTION

A wide range of solute carrier (SLC) and ATP binding cassette transporters, expressed on plasma membranes of multiple organs (e.g., liver, kidney, intestine, brain, placenta), play critical roles in the absorption, distribution, metabolism, and renal and biliary excretion of a multitude of drugs (*1*). Transporter-mediated drug absorption and disposition are saturable and can be modulated by other drugs, potentially leading to drug-drug interactions (DDIs) (*2*). Current DDI guidance documents by the International Council for Harmonisation of Technical Requirements for Pharmaceuticals for Human Use (ICH M12 guidance), U.S. Food and Drug Administration, and European Medicines Agency recommend that drug developers evaluate the inhibition potential of a new chemical entity (NCE) for a range of transporters such as breast cancer resistance protein (BCRP), multidrug and toxin extrusion protein 1 (MATE1), organic anion transporter 1 (OAT1), OAT3, organic anion transporting polypeptides 1B1 (OATP1B1), OATP1B3, organic cation transporter 2 (OCT2), and P-glycoprotein (P-gp) (*3–5*). These guidance documents suggest conducting *in vitro* transporter inhibition assays using probe substrates, followed by DDI predictions using basic static models. However, conventional DDI predictions using these models are associated with a high rate of false-positives, often triggering unnecessary clinical DDI studies, leading to high cost of drug development and potential drug toxicity risks (*6*). Therefore, the ICH M12 guidance suggests alternative approaches, such as mechanistic static models, physiologically-based pharmacokinetic models, and/or endogenous biomarkers, to predict potential DDIs in the clinic, eliminating the need for dedicated DDI studies with exogenous probe drug substrates (*3*). For example, coproporphyrin I (CPI) has increasingly been used as an OATP1B1 biomarker to predict potential OATP1B1-mediated DDIs in the clinic (*7–9*), highlighting the promise of endogenous compounds as transporter biomarkers to assess the DDI potential of a NCE during Phase I single ascending dose (SAD) or multiple ascending dose (MAD) studies. Endogenous substrates of other hepatic and renal transporters include isobutyryl carnitine (OCT1), N1-methylnicotinamide and creatinine (OCT2/MATE), and pyridoxic acid (OAT1/3) (*1*).

Renal SLC transporters, such as OAT1 and 3, are located on the basolateral membrane of proximal tubule cells and work in tandem with apically expressed multidrug resistance-associated protein (MRP) 2 and MRP4 to eliminate drugs (e.g., furosemide, famotidine, methotrexate, and adefovir) and endogenous metabolites (e.g., uremic toxins, tryptophan metabolites, and steroids) (*10*). OAT1/3 expression is affected by factors including age, sex, genotype, and disease state, in addition to being susceptible to DDIs (*11, 12*). In addition to pyridoxic acid, kynurenic acid, homovanillic acid, taurine, glycochenodeoxycholate-3-sulphate (GCDCA-S), and 6β-hydroxycortisol (6βHC) are endogenous substrates of OAT1/3 and have been proposed as biomarkers to predict OAT1/3-mediated DDIs in preclinical species and humans (*13–18*). However, these biomarkers are often confounded by the involvement of other drug metabolizing enzyme and transporter (DMET) proteins, high interindividual variability, reabsorption, and/or non-significant changes in plasma concentrations upon OAT1/3 inhibition (*14, 16, 19*). Thus, there is an unmet need to identify and qualify robust biomarkers of OAT1/3 that can be used to predict not only transporter-mediated DDIs but also interindividual variability in baseline transport function among healthy and specific human populations. Due to the increased sensitivity of liquid chromatography-tandem mass spectrometry (LC-MS)-based metabolomics approaches, biomarker discovery has become more straightforward (*20, 21*). Untargeted metabolomics is particularly powerful for biomarker discovery and quantification as it involves simultaneous unbiased analysis of many small molecules such as endogenous metabolites, xenobiotics, and drug metabolites in biological matrices (*22*).

Discovery of transporter biomarkers can be achieved by coupling metabolomics with *in vitro* models, *in vivo* preclinical models, and/or clinical studies (*23*). *In vitro* cell culture models can be used to assess transporter-mediated uptake of endogenous metabolites by incubating the transporter over-expressing cells with purified standards or biological matrices containing the endogenous metabolites (*23*). This approach can be combined with preclinical models and clinical studies for biomarker validation. Preclinical models such as knockout mice and/or inhibitor administration to wild type animals have been used to study the effects of deletion or transient inhibition of the transporter of interest on the plasma and/or urine metabolome, respectively (*24–27*). Metabolomics analysis of plasma from *Oat1* or *Oat3* knockout mice and *Oat1*/*3* double-knockout rats revealed that concentrations of gut microbiome-generated metabolites, glucuronide metabolites, tryptophan metabolites, uremic toxins, and steroids are altered relative to wild type animals (*22, 24–31*). Shen et al. reported that probenecid-mediated inhibition of renal OAT1/3 significantly increased plasma concentrations and decreased the renal clearance (CL_R_) of pyridoxic acid and homovanillic acid in cynomolgus monkeys (*15*). Using adult healthy rats as a model for Oat1/3 inhibition, we identified 21 features as putative biomarkers of Oat1/3 (*32*). Clinical studies can also be used to identify transporter biomarkers through genome-wide association studies (GWAS), pharmacogenetic studies, and DDI studies. Yee et al. utilized publicly available GWAS data, along with DDI studies and *in vitro* assays, to identify bile acids and fatty acid dicarboxylates as putative biomarkers of hepatic OATP1B1 (*33*). Similarly, various clinical metabolomics analyses during the last decade included the effects of probenecid-mediated inhibition of OAT1/3 transport activity on plasma and urine concentrations of endogenous metabolites (*14, 16, 34*). Although these extensive studies identified new biomarkers of OAT1/3 function, they are often limited by the need for follow-up *in vitro* confirmatory studies or clinical pharmacokinetic analyses.

We developed a novel approach, termed metabolomics-based DMET biomarker discovery (MDBD), to identify and qualify (*via in vitro* mechanistic confirmation and clinical pharmacokinetic validation) clinical biomarkers of OAT1/3 transport function in humans. This multi-step approach (Fig. 1) began with identifying clinical biomarkers using untargeted metabolomics of biological samples obtained from a clinical probenecid-furosemide pharmacokinetic DDI study (n=16). Next, the identified biomarkers were mechanistically confirmed as substrates of OAT1/3 using *in vitro* assays in transporter over-expressing human embryonic kidney (HEK)-293 cells, followed by clinical validation via pharmacokinetic analysis (0-24 h) of plasma and urine from four clinical study participants. The structures of the validated biomarkers can be characterized using a MS/MS fragmentation approach and compared with analytical standards. Select biomarkers discovered through the MDBD approach can be combined into a panel to evaluate alterations in OAT1/3 transport function by various intrinsic (e.g., age, sex, disease state, pregnancy, kidney function) and extrinsic (e.g., drugs, food, natural products) factors. This novel approach can be applied to the discovery of biomarkers for other DMET proteins.

**Figure 1.**
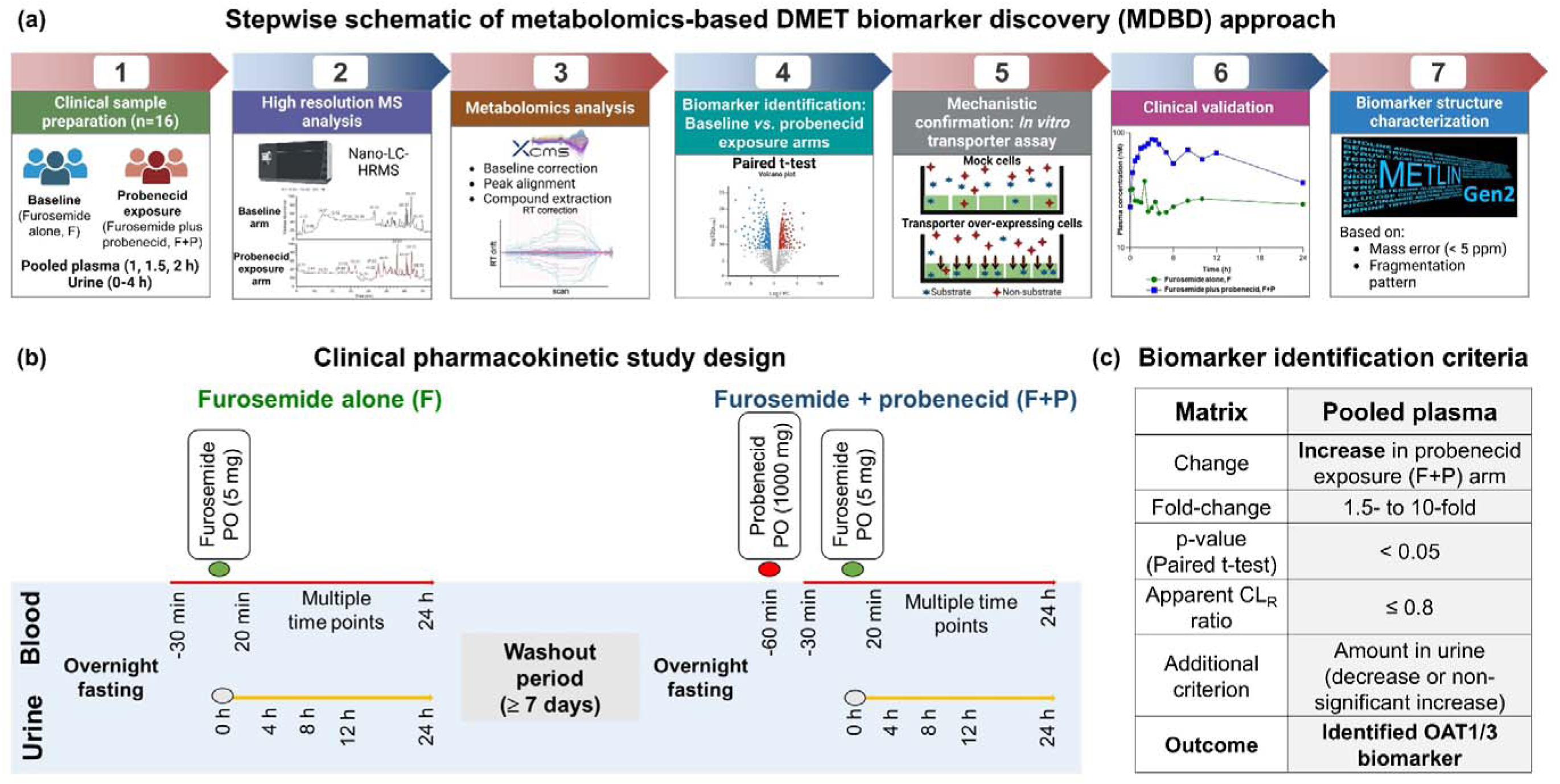
Metabolomics-based DMET biomarker discovery (MDBD) approach. (a) Stepwise workflow of the MDBD approach; (b) Clinical pharmacokinetic drug-drug interaction study design involving 16 healthy adult participants; and (c) Biomarker identification criteria used for the analysis of untargeted metabolomics data of pooled plasma and urine samples.

## RESULTS

### Probenecid coadministration alters the pharmacokinetics of furosemide

All participants completed both arms of the study. Both furosemide and probenecid were well-tolerated by all participants, with no to minimal adverse effects. Probenecid-mediated inhibition of renal OAT1/3 led to a significant increase in plasma concentrations of furosemide at the 40 min, 1 h, 1.5 h, 2 h, 2.5 h, 3 h, 3.5 h, 4 h, 5 h, 6 h, 8 h, 10 h, and 12 h time-points (p-value < 0.05). Inhibition of furosemide renal secretion resulted in a significant increase in average AUC_0-24 h_ (462.4 ± 124.1 ng.h/mL to 854.6 ± 246.1 ng.h/mL), with a significant decrease in the amount excreted unchanged in urine from 0-24 h (A_e, 0-24 h_; 2.6 ± 0.7 mg to 1.7 ± 0.5 mg). This resulted in a decrease in furosemide CL_R_ from 96.1 ± 27.4 mL/min to 33.4 ± 9.2 mL/min (p-value < 0.05) (Fig. 2).

**Figure 2.**
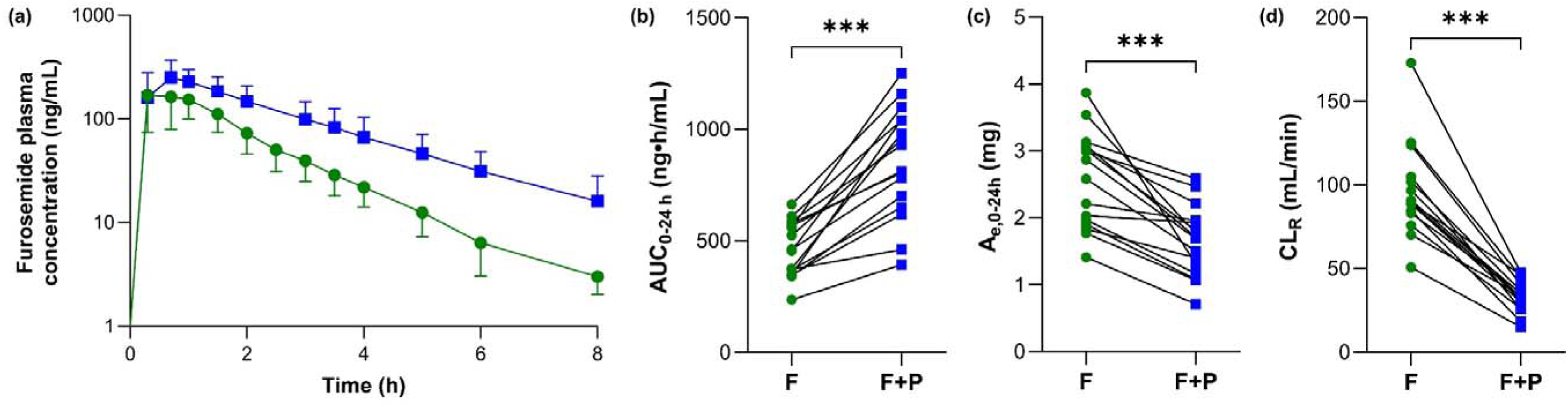
Pharmacokinetic profile of furosemide in the presence and absence of probenecid. (a) Average plasma concentration *versus* time profiles, and individual (b) AUC_0-24 h_, (c) A_e,0-24 h_, and (d) CL_R_ of furosemide in adult healthy participants (n=16) after oral administration of furosemide either alone (green circles, F) or with probenecid (blue squares, F+P). The average data represent geometric means with error bars indicating standard deviation. Pharmacokinetic endpoints were compared between the baseline and the probenecid exposure arms using the paired t-test. p-value < 0.0001 (***). F = furosemide alone (baseline arm); F+P = furosemide coadministered with probenecid (probenecid exposure arm).

### Metabolomics-based DMET biomarker discovery (MDBD) approach

Plasma and urine samples from the probenecid-furosemide DDI study were used to identify putative biomarkers of OAT1 and OAT3 using the MDBD approach (Fig. 1a-b). We hypothesized that inhibition of renal OATs by probenecid would lead to an increase in plasma concentrations and a corresponding decrease in the CL_R_ of substrates. Based on literature reports (*14, 35*) and preliminary assessment of probenecid exposure, plasma samples (pool of 1, 1.5, and 2 h) and urine samples (0-4 h) representing maximum probenecid concentration (and thus maximum inhibitory potential) were selected for biomarker identification. Upon probenecid exposure, a total of 631 features detected in both positive and negative ionization modes showed significant increases in pooled plasma (1.5- to 10-fold), with 153 features showing a corresponding decrease in apparent CL_R_ by more than 20% (Fig. 1c). Among these, 57 features were mechanistically confirmed as substrates of OAT1 and/or OAT3 via *in vitro* transporter assays using transporter over-expressing cells. Finally, 23 features were clinically validated as biomarkers of OAT1/3 through detailed pharmacokinetic analysis (0-24 h) of plasma and urine samples from four study participants. Biomarkers with consistent pharmacokinetic profiles from both the baseline and probenecid exposure arms were selected. The structure of one representative OAT1/3 biomarker (*m/z* 162.0550) was characterized using a robust approach involving MS/MS fragmentation combined with spiking the plasma with authentic reference standard.

### Probenecid alters the plasma and urine metabolome

Principal component analysis of the untargeted metabolomics data of pooled plasma revealed that individual participants were distinguishable between the baseline and probenecid exposure arms, suggesting a significant physiological effect of probenecid treatment (Supplementary Fig. 1). Untargeted analysis of pooled plasma samples in positive ionization mode yielded 7,962 features (unique *m/z* values), of which 7511 were common between the two arms, while 84 and 367 features were unique to the baseline and probenecid exposure arms, respectively (Fig. 3). Similarly, in negative ionization mode, 5,043 features from pooled plasma samples were common between the baseline and probenecid exposure arms (Supplementary Fig. 2). In the 0-4 h urine samples, 11,562 and 6,210 features were common between the two arms in positive and negative ionization modes, respectively (Supplementary Fig. 3). Statistical analysis of pooled plasma revealed that 907 and 743 features were significantly increased in the probenecid exposure arm in positive and negative modes, respectively (Fig. 3, Supplementary Fig. 2, Supplementary Datafile 1a). Additionally, 474 and 609 features were significantly decreased in the probenecid exposure arm compared to the baseline arm in positive and negative modes, respectively (Fig. 3, Supplementary Fig. 2, Supplementary Datafile 1b). Regarding the urine samples, 2,842 features (1,830 in positive mode and 1,012 in negative mode) were significantly increased, while 1,910 features (659 in positive mode and 1,251 in negative mode) were significantly decreased upon probenecid exposure (Supplementary Fig. 3).

**Figure 3.**
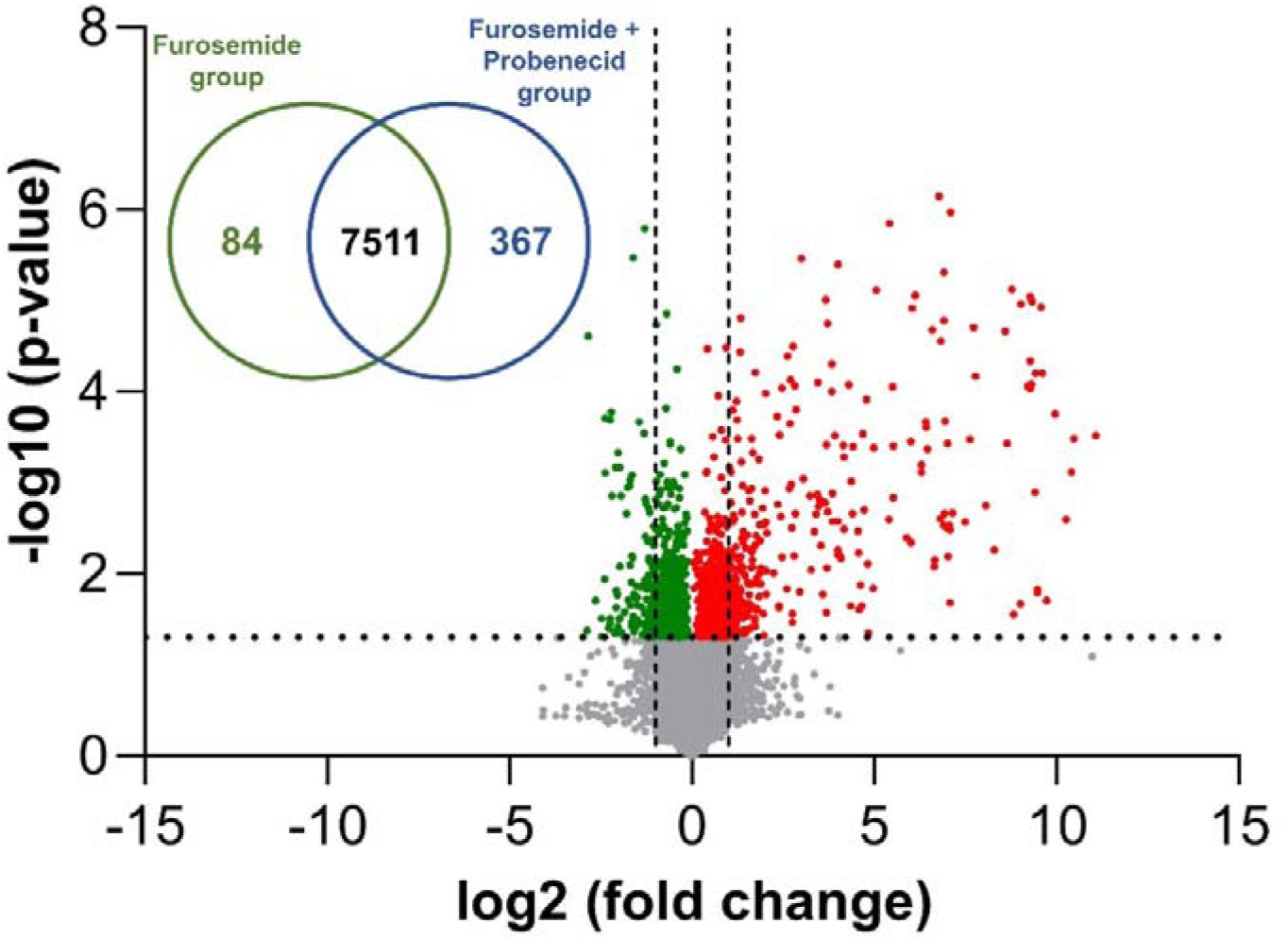
Comparison of *m/z* features detected in pooled plasma (1, 1.5, 2 h) in positive ionization mode in the baseline and probenecid exposure arms. Venn diagram showing the total number of unique and common features detected in the baseline and probenecid exposure arms. Volcano plot of the detected features, with dots representing *m/*z features that were either increased or decreased in pooled plasma in the presence of probenecid. Red and green dots indicate *m/z* features that were significantly higher or lower (p-value < 0.05) in pooled plasma on probenecid exposure, respectively. The horizontal and vertical lines represent the p-value (< 0.05) and fold change (> or < 2-fold), respectively.

### Identification of putative OAT1/3 biomarkers

Analysis of pooled plasma in positive mode identified 377 features that were significantly increased (by 1.5- to 10-fold) in the probenecid exposure arm (Supplementary Datafile 1a). In negative ionization mode, 254 features were significantly increased (by 1.5- to 10-fold) in pooled plasma with probenecid exposure compared to the baseline arm (Supplementary Datafile 1a). For example, the feature corresponding to an *m/z* of 162.0550 showed a 2.1-fold higher pooled plasma concentration in the probenecid exposure arm compared to the baseline arm (p-value > 0.05), with a corresponding 45% decrease in apparent CL_R_. In contrast, examples such as a feature with *m/z* 195.1020 exhibited a greater increase in pooled plasma concentration (in the probenecid exposure arm, 5-fold, p-value < 0.05), but with an increase in apparent CL_R_ (∼25% increase), leading to their exclusion from the list of putative OAT1/3 biomarkers due to lack of adherence to the selection criteria (Fig. 1c). Thus, screening of the features based on the selection criteria (1.5- to 10-fold increase in the pooled plasma concentration in the probenecid exposure arm with p-value < 0.05 and a ≥20% decrease in apparent CL_R_) yielded a list of 153 features (88 and 65 in the positive and negative ionization modes, respectively) identified as putative OAT1/3 biomarkers, with *m/z* values ranging from 110 to 510. Based on METLIN-predicted names, the identified biomarkers belonged to metabolic categories including steroids, peptides, glucuronide conjugates, fatty acids, and tryptophan and tyrosine metabolites.

### *In vitro* mechanistic confirmation of OAT1/3 biomarkers

The identified putative OAT1/3 biomarkers were mechanistically confirmed as OAT1/3 substrates based on the untargeted analysis of *in vitro* transporter uptake assay samples. Untargeted metabolomics analysis yielded >5,500 features, of which ∼5,000 features were shared between the mock and OAT1 over-expressing cells or between mock and OAT3 over-expressing cells (Fig. 4a, Supplementary Fig. 4a). Features showing >1.5-fold higher uptake in OAT1 or OAT3 over-expressing cells were attributed to OAT1 or OAT3 transport (Fig. 4b-c, Supplementary Fig. 4b-c, Supplementary Datafiles 2-3). Compared to mock cells, furosemide (positive control) uptake was 1.8-fold and 1.4-fold higher in OAT1 and OAT3 over-expressing cells, respectively (Supplementary Fig. 5, Supplementary Fig. 6). Therefore, a cutoff value of >1.5-fold uptake in transporter-overexpressing cells was chosen for substrate screening.

**Figure 4.**
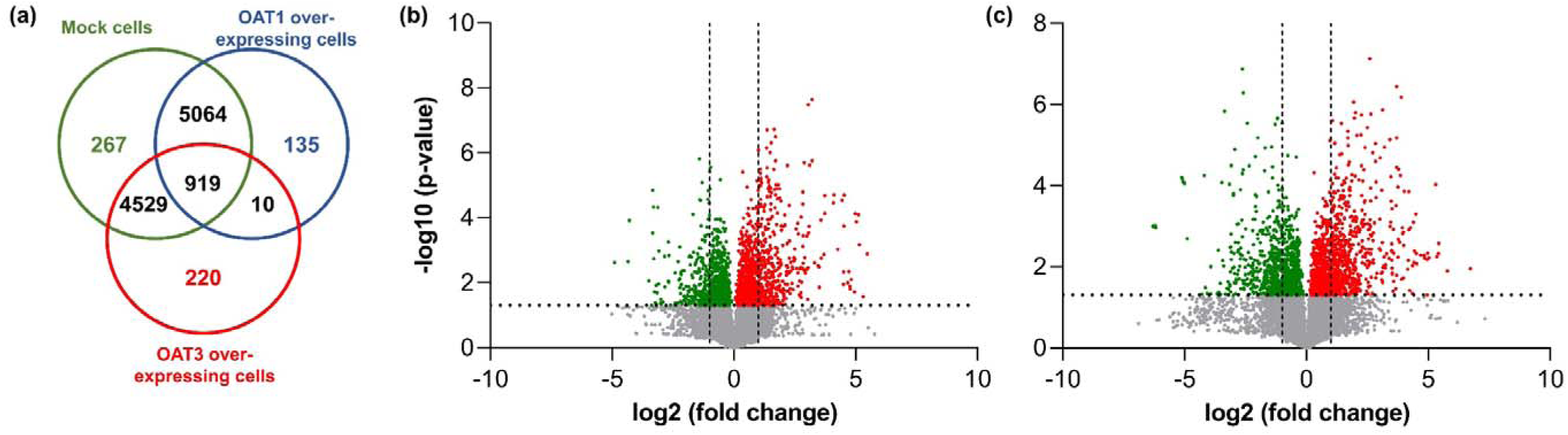
Comparison of *m/z* features concentrated in mock and OAT1 or OAT3 over-expressing cells following *in vitro* uptake assays in positive ionization mode. (a) Venn diagram showing the total number of unique and common features concentrated in mock cells, OAT1 over-expressing cells, and OAT3 over-expressing cells. Volcano plots of the concentrated features, with dots representing *m/*z features that showed either higher or lower uptake in (b) OAT1 over-expressing cells or (c) OAT3 over-expressing cells, compared to mock cells. Red and green dots indicate features with significantly higher or lower uptake (p-value < 0.05) in transporter over-expressing cells, respectively. The horizontal and vertical lines represent the p-value (< 0.05) and fold change (> or < 2-fold), respectively.

The cellular uptake of 1,073 features (500 and 573 in the positive and negative modes, respectively) was significantly increased (>1.5-fold) in OAT1 over-expressing cells compared to mock cells (Supplementary Datafile 2). The *m/z* values of these features ranged from 70 to 616, with a maximum uptake of 44-fold for a feature with an *m/z* of 261.0210. Similarly, 1,523 features (733 in positive mode and 790 in negative mode) showed significantly higher uptake in OAT3 over-expressing cells compared to mock cells (>1.5-fold, Supplementary Datafile 3). The uptake of a feature with an *m/z* of 351.1010 was 105-fold higher in OAT3 over-expressing cells compared to mock cells. Overall, the *m/z* values of OAT3 substrates ranged from 70 to 747. Comparing the *m/z* values of putative OAT1/3 biomarkers identified in clinical samples with the *in vitro* uptake data confirmed 57 features as substrates of OAT1, OAT3, or both (Supplementary Datafile 4). Potential confounding effects of other transporters were also assessed by determining the uptake of these features in OAT2, OATP1B1, and OATP1B3 over-expressing cells.

### Pharmacokinetic analysis of mechanistically confirmed OAT1/3 biomarkers

For the 57 features shortlisted based on *in vitro* data, pharmacokinetic analysis of plasma and urine samples from 0-24 h was conducted for four of the clinical study participants. These participants were selected based on maximum (n=2) and minimum (n=2) changes in furosemide pooled plasma concentration in the presence of probenecid. Based on the pharmacokinetic profiles, as well as AUC and CL_R_ ratios, 23 features were selected as clinically validated biomarkers of OAT1/3. For the 23 validated biomarkers (*m/z* values: 160 to 430), the average pooled plasma concentrations increased by 1.5- to 3.3-fold in the presence of probenecid, with a 20-70% decrease in apparent CL_R_ values in the probenecid exposure arm compared to the baseline arm (Fig. 5, Supplementary Fig. 7). For a feature with an *m/z* value of 162.0550, the average pooled plasma concentration was 2.1 ± 1.1-fold higher in the probenecid arm compared to the baseline arm and showed a ∼50% decrease in average apparent CL_R_ values (Fig. 5, Supplementary Fig. 7). The uptake of this biomarker was 3-, 2.1-, and 2.5-fold higher in OAT1, OAT3, and OAT2 over-expressing cells, respectively, compared to mock cells, suggesting that it is a substrate for these transporters (Supplementary Figs. 5 and 6, Supplementary Datafile 5). Analysis of the 0-24 h pharmacokinetic data for this feature in the four participants revealed that baseline variability in plasma concentration was <30%, with minimal fluctuations except for increased concentrations at the 24 h time-point. Administration of probenecid significantly increased plasma concentrations of this biomarker, with t_max_ ranging from 3-5 h, followed by a decrease as probenecid was eliminated from the body (Fig. 6, Supplementary Fig. 8). The AUC and CL_R_ ratios were 2.2 ± 0.4 and 0.4 ± 0.2, respectively (n=4, Table 1, Supplementary Datafile 6). In contrast, a feature with an *m/z* of 245.0921 (average pooled plasma fold change = 2.9; apparent CL_R_ = 0.4) showed significantly higher uptake in only OAT1 (2.3-fold) and OAT3 (4.1-fold) over-expressing cells. This biomarker had low baseline variability (<25%), but variability in the probenecid arm was high (AUC ratio = 3.3 ± 3.4; CL_R_ ratio = 0.4 ± 0.1). Features with *m/z* values of 310.2013, 188.0928, 269.1510, and 427.1974 were selective substrates of OAT3, with 1.9-, 1.5-, 4.5-, and 1.6-fold higher uptake in OAT3 over-expressing cells compared to mock cells (p-value < 0.05). Although features with *m/z* 251.1278, 269.1384, and 319.1904 were more selective for OAT1 compared to OAT3, these biomarkers were also substrates of one or more other transporters (OAT2 and OATP1B3). The feature with *m/z* 251.1278 is potentially a water loss product of *m/z* 269.1384.

**Figure 5.**
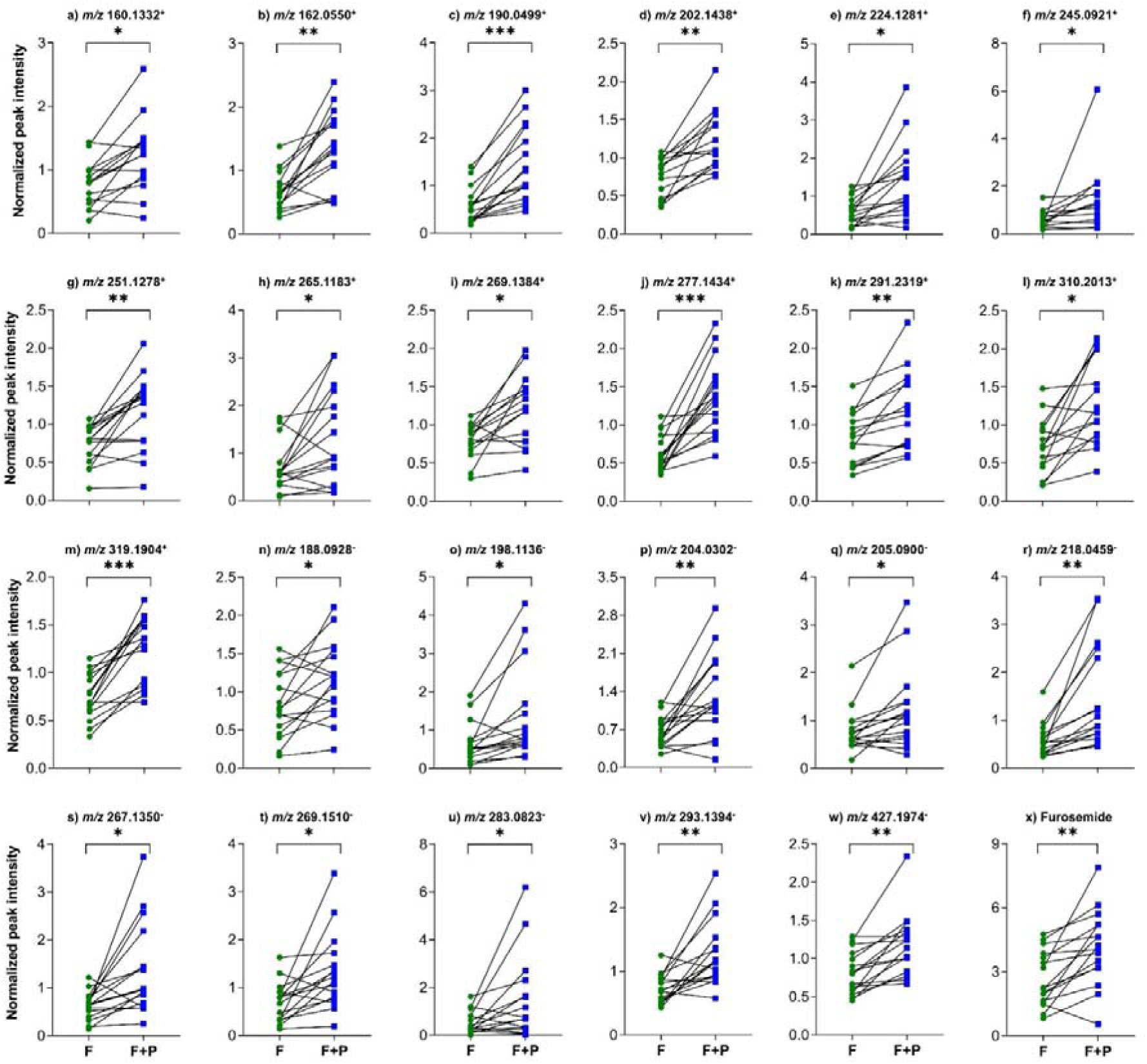
Comparison of the pooled plasma concentrations of validated OAT1/3 biomarkers and furosemide in the baseline and probenecid exposure arms. Line and symbol plots of validated OAT1/3 biomarkers (a-w) and furosemide (x) with normalized peak intensity on the y-axis representing pooled plasma concentration in healthy adult participants (n=16) in baseline (green circles, F) and probenecid exposure (blue squares, F+P) arms. Normalized peak intensities in pooled plasma were compared between the baseline and probenecid exposure arms using the paired t-test. p-value < 0.05 (*), < 0.001 (**), and < 0.0001 (***). F = furosemide alone (baseline arm); F+P = furosemide coadministered with probenecid (probenecid exposure arm); *m/z*: Mass-to-charge ratio with positive and negative sign in superscript representing the ionization mode.

**Figure 6.**
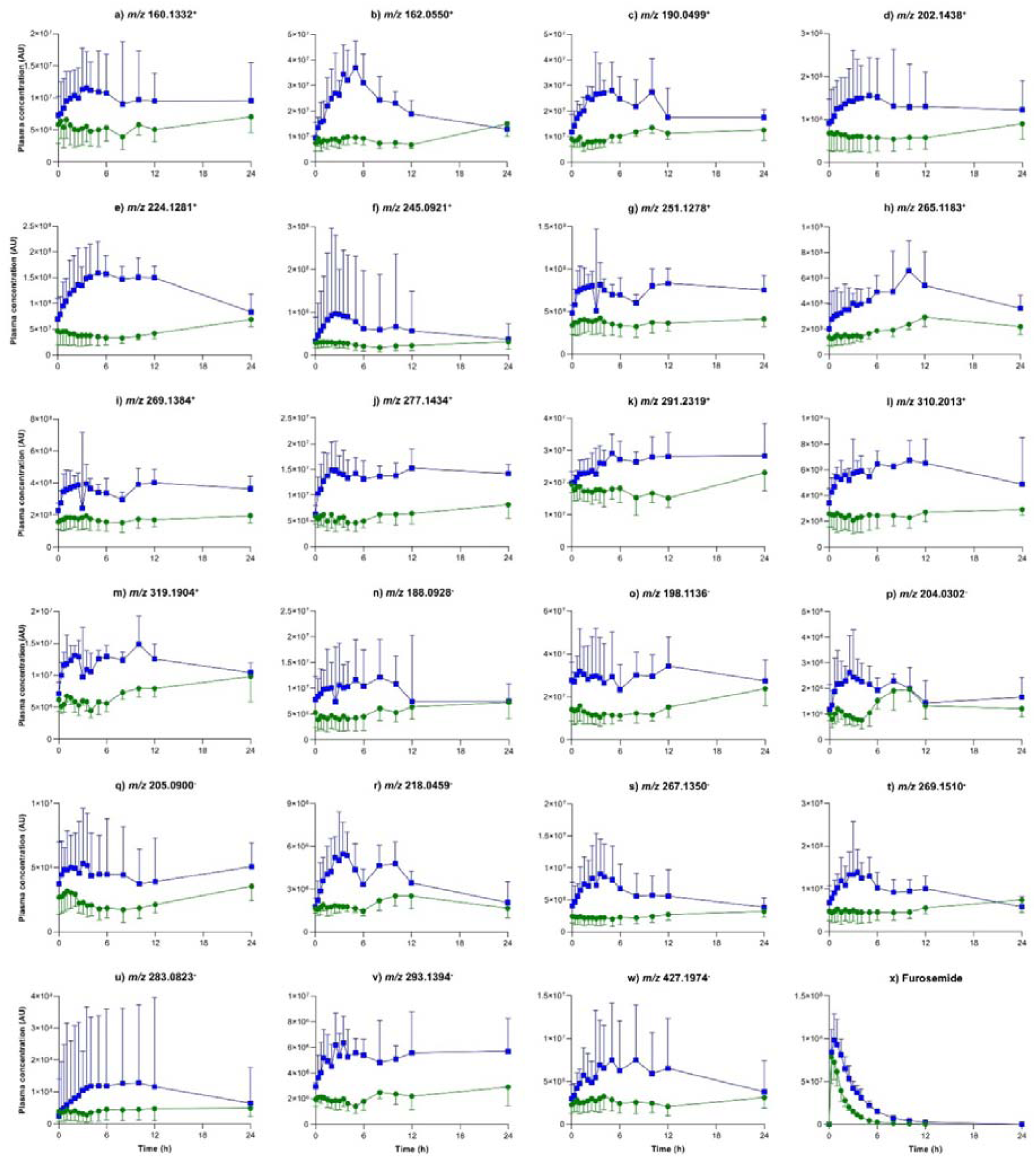
Pharmacokinetic profiles of the promising OAT1/3 biomarkers and furosemide. Average plasma concentration (peak intensities) *versus* time profiles of validated OAT1/3 biomarkers (a-w) and furosemide (x) in healthy adult participants (n=4) in baseline (green circles, F) and probenecid exposure (blue squares, F+P) arms. The average data represent geometric means with error bars indicating standard deviation. F = furosemide alone (baseline arm); F+P = furosemide coadministered with probenecid (probenecid exposure arm); AU = Arbitrary units; *m/z*: Mass-to-charge ratio with positive and negative sign in superscript representing the ionization mode.

**Table 1.**
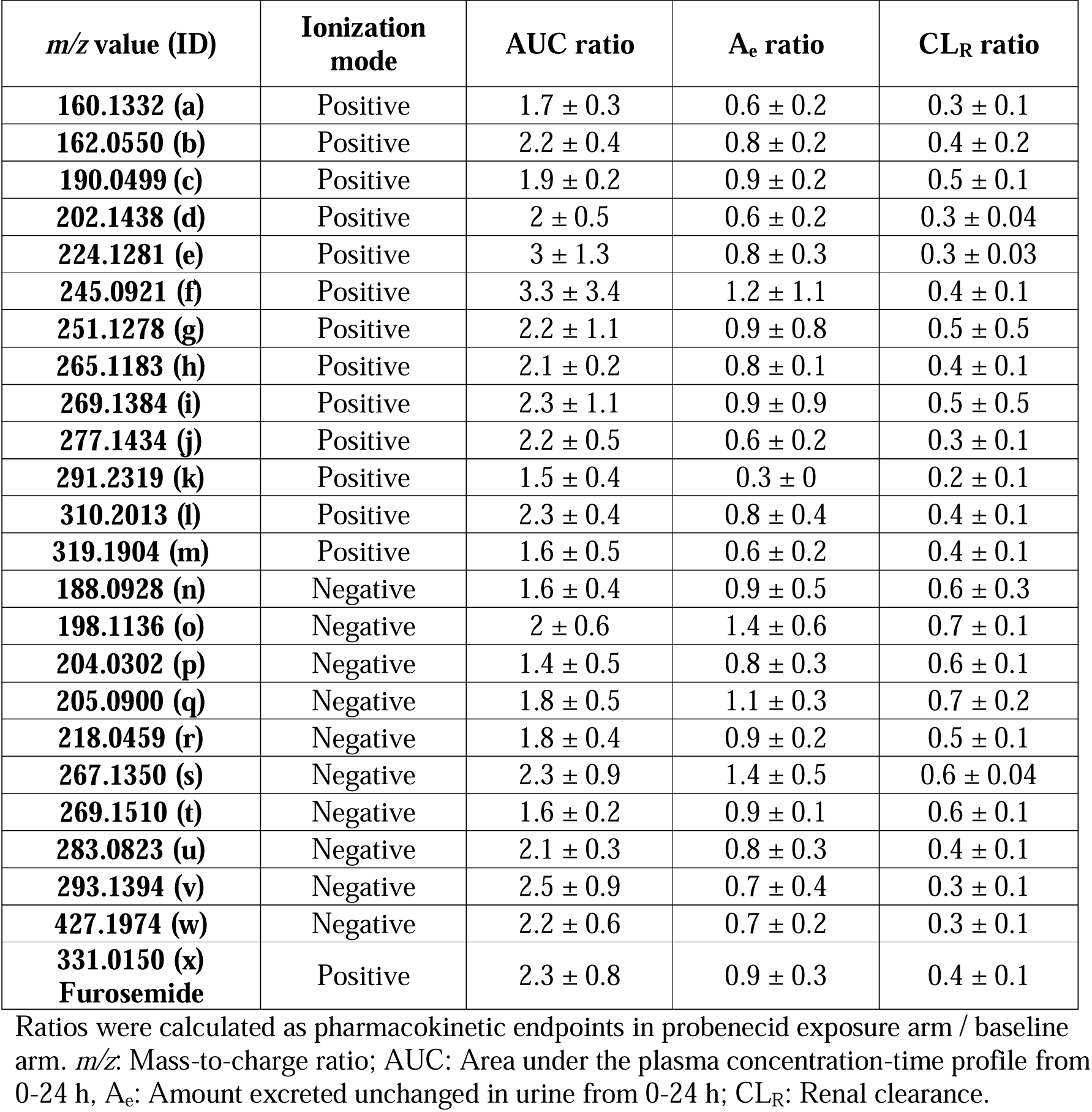
Average fold change (± standard deviation) in pharmacokinetic endpoints of validated OAT1/3 biomarkers and furosemide on probenecid-mediated inhibition of renal OATs in healthy adult participants (n=4).

| *m/z* value (ID) | Ionization mode | AUC ratio | A<sub>e</sub> ratio | CL<sub>R</sub> ratio |
| --- | --- | --- | --- | --- |
| **160.1332 (a)** | Positive | 1.7 ± 0.3 | 0.6 ± 0.2 | 0.3 ± 0.1 |
| **162.0550 (b)** | Positive | 2.2 ± 0.4 | 0.8 ± 0.2 | 0.4 ± 0.2 |
| **190.0499 (c)** | Positive | 1.9 ± 0.2 | 0.9 ± 0.2 | 0.5 ± 0.1 |
| **202.1438 (d)** | Positive | 2 ± 0.5 | 0.6 ± 0.2 | 0.3 ± 0.04 |
| **224.1281 (e)** | Positive | 3 ± 1.3 | 0.8 ± 0.3 | 0.3 ± 0.03 |
| **245.0921 (f)** | Positive | 3.3 ± 3.4 | 1.2 ± 1.1 | 0.4 ± 0.1 |
| **251.1278 (g)** | Positive | 2.2 ± 1.1 | 0.9 ± 0.8 | 0.5 ± 0.5 |
| **265.1183 (h)** | Positive | 2.1 ± 0.2 | 0.8 ± 0.1 | 0.4 ± 0.1 |
| **269.1384 (i)** | Positive | 2.3 ± 1.1 | 0.9 ± 0.9 | 0.5 ± 0.5 |
| **277.1434 (j)** | Positive | 2.2 ± 0.5 | 0.6 ± 0.2 | 0.3 ± 0.1 |
| **291.2319 (k)** | Positive | 1.5 ± 0.4 | 0.3 ± 0 | 0.2 ± 0.1 |
| **310.2013 (l)** | Positive | 2.3 ± 0.4 | 0.8 ± 0.4 | 0.4 ± 0.1 |
| **319.1904 (m)** | Positive | 1.6 ± 0.5 | 0.6 ± 0.2 | 0.4 ± 0.1 |
| **188.0928 (n)** | Negative | 1.6 ± 0.4 | 0.9 ± 0.5 | 0.6 ± 0.3 |
| **198.1136 (o)** | Negative | 2 ± 0.6 | 1.4 ± 0.6 | 0.7 ± 0.1 |
| **204.0302 (p)** | Negative | 1.4 ± 0.5 | 0.8 ± 0.3 | 0.6 ± 0.1 |
| **205.0900 (q)** | Negative | 1.8 ± 0.5 | 1.1 ± 0.3 | 0.7 ± 0.2 |
| **218.0459 (r)** | Negative | 1.8 ± 0.4 | 0.9 ± 0.2 | 0.5 ± 0.1 |
| **267.1350 (s)** | Negative | 2.3 ± 0.9 | 1.4 ± 0.5 | 0.6 ± 0.04 |
| **269.1510 (t)** | Negative | 1.6 ± 0.2 | 0.9 ± 0.1 | 0.6 ± 0.1 |
| **283.0823 (u)** | Negative | 2.1 ± 0.3 | 0.8 ± 0.3 | 0.4 ± 0.1 |
| **293.1394 (v)** | Negative | 2.5 ± 0.9 | 0.7 ± 0.4 | 0.3 ± 0.1 |
| **427.1974 (w)** | Negative | 2.2 ± 0.6 | 0.7 ± 0.2 | 0.3 ± 0.1 |
| **331.0150 (x)**<br>**Furosemide** | Positive | 2.3 ± 0.8 | 0.9 ± 0.3 | 0.4 ± 0.1 |

Ratios were calculated as pharmacokinetic endpoints in probenecid exposure arm / baseline arm. *m/z*: Mass-to-charge ratio; AUC: Area under the plasma concentration-time profile from 0-24 h, A<sub>e</sub>: Amount excreted unchanged in urine from 0-24 h; CL<sub>R</sub>: Renal clearance.

A feature with an *m/z* of 218.0459 exhibited a 2.8-fold increase in pooled plasma concentration in the presence of probenecid and had 20- to 45-fold higher uptake in OAT1, OAT2, or OAT3 over-expressing cells. However, 0-24 h pharmacokinetic analysis revealed double peaks in both the baseline and probenecid exposure arms for this biomarker, suggesting diurnal variation or other contributing factors (Fig. 6). Similarly, a feature with an *m/z* value of 204.0302 showed double peaks in both the baseline and probenecid exposure arms (Fig. 6).

### Structure characterization of OAT1/3 biomarkers: Case example of *m/z* 162.0550

The feature with *m/*z 162.0550 is a sensitive OAT1/3 biomarker that showed a 2.2-fold increase in AUC ratio with a corresponding decrease in CL_R_ and minimal baseline variability. Therefore, we characterized the structure of this biomarker. MS/MS-based fragmentation patterns were leveraged for structure characterization. METLIN predicted the following 10 putative names: Indole-2-carboxylic acid, indole-3-carboxylic acid, 2,8-dihydroxyquinoline, 3,4-dihydroxyquinoline, 4,6-dihydroxyquinoline, 4,8-dihydroxyquinoline, 3-hydroxy-1H-quinolin-4-one, 4-hydroxy-1H-quinolin-2-one, 3-formyl-6-hydroxyindole, and 1,5-dihydroxy-isoquinoline (Fig. 7a). MS/MS fragmentation of the feature revealed two major fragments: *m/z* 144.0444 (water loss) and *m/*z 118.0651 (carbon dioxide loss). Based on the structures of METLIN-predicted compounds, the losses of water and carbon dioxide are only possible for indole-2-carboxylic acid (I2CA) and indole-3-carboxylic acid (I3CA) (Fig. 7b). Analytical standards of these two compounds were procured to confirm the feature identity by matching LC retention times and MS/MS fragments. Although both I2CA and I3CA standards produced the neutral losses observed in the biomarker, the retention times of the LC peaks for I2CA and I3CA standards were 18.4 and 16 min, respectively, compared to the retention time of 14.4 min for the validated biomarker (Fig. 7b-c). This observation suggests that the biomarker is neither of these compounds.

**Figure 7.**
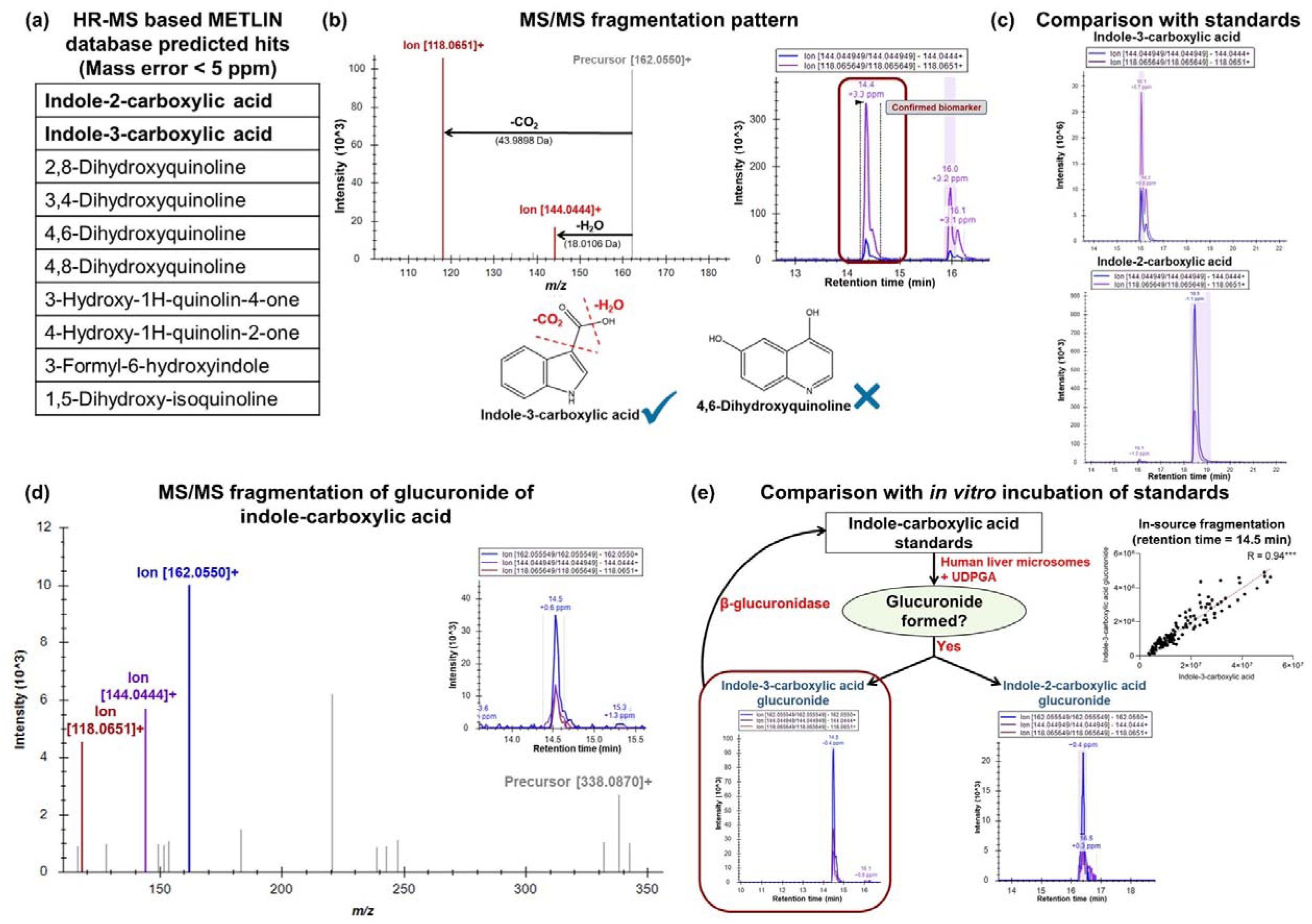
Stepwise approach for structure elucidation of OAT1/3 biomarker, *m/z* 162.0550. (a) Known compounds with the high-resolution mass spectrometry (HR-MS) predicted *m/z* value of 162.0550 (mass error < 5 ppm) in METLIN database search. (b) MS/MS fragmentation pattern of the OAT1/3 biomarker, *m/z* value of 162.0550, along with the chromatographic peak of the feature fragments. Fragments with *m/z* values of 144.0444 and 118.0651 result from water and carbon dioxide loss, respectively. Water and carbon dioxide losses are only possible from indole-carboxylic acid (red dashed lines). (c) Chromatograms of the individual fragments of the indole-carboxylic acid standards. (d) MS/MS fragmentation pattern of the glucuronide of indole-carboxylic acid. The inset shows chromatograms of individual fragments. (e) Synthesis of indole-carboxylic acid glucuronide from indole-carboxylic acid standards using human liver microsomes and correlation of indole-3-carboxylic acid glucuronide (I3CAG) with the in-source fragmentation product, I3CA at LC retention time of 14.5 min. I3CAG = Indole-3-carboxylic acid glucuronide; *m/z* = Mass-to-charge ratio; METLIN = Metabolite and Chemical Entity Database; and UDPGA = uridine diphosphate glucuronic acid.

Because the biomarker and the analytical standards showed similar mass fragmentation profiles, we hypothesized that the biomarker is a derivative of either I2CA or I3CA. The more hydrophilic nature of the biomarker compared to the standards (shorter retention time) suggested that the biomarker could potentially be a conjugate of one of these compounds. Mass fragmentation of conjugate masses at the biomarker LC retention time (14.4 min) revealed that the biomarker is a glucuronide of indole-carboxylic acid (Fig. 7d). Subsequently, incubation of the analytical standards with human liver microsomes produced the glucuronides of both I2CA and I3CA, with similar MS/MS fragmentation patterns. However, the retention time of the I3CA glucuronide (I3CAG) (14.5 min) matched closely with that of the biomarker (Fig. 7e), suggesting that the feature with *m/z* 162.0550 is a surrogate of the original biomarker, I3CAG, and is formed by in-source loss of glucuronic acid. This conclusion is supported by a strong correlation (r > 0.9, p-value < 0.0001) between the peak intensities of I3CAG at 14.5 min retention time and the peak intensities of the in-source fragmented product, I3CA, at the same retention time (Fig. 7e). This observation suggests that the in-source fragmentation is linear, and that I3CA can be used as a surrogate to measure the concentration of I3CAG. The literature suggests that the glucuronide of I3CA is an acyl glucuronide, rather than an N-glucuronide (*36*). We confirmed this by using a hydroxylamine derivatization approach, wherein hydroxylamine will displace the acyl glucuronide to yield the hydroxamic acid derivative of the compound, while not reacting with an N-glucuronide (*37*). The reaction of hydroxylamine with processed urine samples yielded indole-3-hydroxamic acid, which was absent in urine untreated with hydroxylamine, confirming that I3CAG is an acyl glucuronide (Supplementary Fig. 9). Further, assessment of I3CAG glucuronidation by 13 recombinant UGT revealed that I3CA is a substrate of UGT1A6, UGT1A7 and UGT1A9 (Supplementary Fig. 10).

If plasma concentration of a biomarker reflects OAT1/3 activity (where OAT-mediated renal secretion is the rate-limiting step), a higher plasma concentration would indicate lower OAT1/3 activity. Reduced transport activity would decrease the fraction of the substrate drug transported by OAT1/3, thus lowering both the DDI risk and the AUC ratio of the substrate drug. The pre-dose (0 min) plasma concentration of I3CAG showed a significant negative correlation with the pooled plasma fold change of furosemide (R > 0.5, p-value < 0.05), suggesting that pre-dose plasma levels of I3CAG could potentially be used to estimate the OAT1/3-mediated DDI risk for substrate drugs (Supplementary Fig. 11).

### Utility of a panel of the top six OAT1/3 biomarkers for prediction of interindividual variability in probenecid-mediated changes in furosemide plasma concentration

The six most promising biomarkers were selected for a panel based on their low intraindividual (< 50%) and interindividual (< 30%) variability in baseline plasma concentrations, lack of diurnal variation, AUC ratio (≥ 2-fold), and CL_R_ ratio (≤ 0.6). Biomarkers chosen for the panel were features with *m/*z values of 162.0550, 202.1438, 277.1434, 310.2013, 267.1350, and 427.1974. The fold change in furosemide concentrations in pooled plasma (n=16) significantly correlated with average fold change in biomarker panel (r = 0.5, p-value < 0.05) (Supplementary Fig. 12), highlighting that OAT1/3-mediated interindividual variability in the probenecid-furosemide interaction can be predicted by a panel of OAT1/3 biomarkers.

## DISCUSSION

The metabolome offers a unique advantage over the transcriptome and proteome by depicting the phenotype of a biological process (*38*). For instance, metabolomics can be used to measure both steady-state concentrations of metabolites and dynamic response of metabolites to intrinsic or extrinsic stimuli, allowing for the investigation of perturbations in biological processes and the discovery of biomarkers related to these dynamic changes (*39*). Although nuclear magnetic resonance (NMR) can provide comprehensive structural information and absolute quantification of metabolic biomarkers, LC-MS has gained traction during the past two decades due to its superior sensitivity, improved metabolome coverage, and high-throughput nature (*20, 21*). LC-MS-based metabolomics can be carried out using either targeted or untargeted approaches; the targeted analysis focuses on known metabolites, while the untargeted approach quantifies a large number of unspecified analytes present in a biological sample (*40*). Therefore, in this study, we leveraged untargeted metabolomics for biomarker discovery as well as quantitative metabolite analysis to investigate the effect of perturbed renal OAT activity on plasma and urine metabolomes. We developed a novel and integrated metabolomics workflow for the identification and qualification of DMET biomarkers for prospective assessment of clinical DDI potential and interindividual variability using renal OAT1 and OAT3 as case examples.

We first conducted a pharmacokinetic DDI study in healthy adults, who were administered OAT substrate, furosemide alone (baseline arm) or with an OAT1/3 inhibitor (probenecid exposure arm). Plasma and urine samples representing peak probenecid-mediated inhibition were used for untargeted metabolomics analysis and identification of putative biomarkers of renal OATs. LC-MS data acquisition was conducted in data-independent (DIA) mode, which provided greater sensitivity for quantifying low abundance metabolites. Inhibition of OAT1/3 leads to an increase in plasma concentrations of OAT1/3 substrates. However, since plasma metabolite concentrations can also be perturbed by off-target effects of probenecid, we analysed both plasma and urine concentrations of the metabolomic features and determined the apparent CL_R_ values. Furthermore, the putative biomarkers identified in the clinical study were confirmed as OAT1/3 substrates by conducting *in vitro* uptake assays in transporter over-expressing cells. Using urine, we first conducted transporter uptake assays to assess putative substrates of OAT1 and OAT3. To establish the selectivity of the putative transporter substrates, they were also assessed for OAT2, OATP1B1 and OATP1B3 uptake. While OATPs are expressed exclusively in the liver, OAT2 is expressed in the liver and kidney. OAT2 abundance in the kidney is 10-fold lower compared to OAT1 and OAT3 (*41, 42*). Additionally, the inhibitory potential of probenecid for OAT2 is low, with >45-fold higher half maximal inhibitory concentration (IC_50_) than that for OAT1 and OAT3 (393 µM vs. 8.2 and 5.8 µM) (*15*). Based on the *in vitro* uptake assay results, 57 features were shortlisted, which were then subjected to the final pharmacokinetic analysis screen. The 0-24 h pharmacokinetic analysis yielded 23 features as clinically validated robust biomarkers of OAT1/3.

Use of single or double knockout mice or rats has identified Oat1/3 substrates such as indoxyl sulfate, indole lactic acid, phenyl sulfate, hippuric acid, cysteine, trimethylamine N-oxide, citrulline, acetyl tryptophan, p-cresol sulfate, and kynurenic acid based on the increase in plasma concentrations following gene knockout (*22, 24–31*). While these studies provide a holistic understanding of transporter function, gene knockout can lead to compensatory changes in alternate or rescue pathways that may not align with changes observed during transient inhibition (*25, 43*). Shen et al. conducted a pharmacokinetic DDI study in cynomolgus monkeys using probenecid/furosemide as the inhibitor/substrate pair and measured 233 metabolites in plasma and urine samples, out of which pyridoxic acid and homovanillic acid were identified as OAT1/3 substrates (*15*). We also identified 21 putative biomarkers of rat Oat1/3 using a pharmacokinetic DDI study wherein probenecid was administered as an inhibitor (*32*). While preclinical models can be used to identify biomarkers of DMET proteins, there are significant differences in the renal abundance of OATs between humans and monkeys or humans and rats (*41, 44*). Additionally, physiological differences in the expression of other transporters among these species (*41*) complicates clinical translation of these data. To address the challenge of inter-species differences, clinical studies have been conducted to discover biomarkers of OAT1/3 (*13, 14, 16, 34*). Pyridoxic acid was validated as a clinical biomarker of OAT1/3, as evidenced by a >3-fold increase in plasma concentration and >60% decrease in CL_R_ in healthy adults, upon administration of single or multiple oral doses of probenecid (*14, 16*). Similarly, kynurenic acid was identified as a putative biomarker of OAT1/3 (*13*). Granados et al. performed targeted metabolomics on plasma and urine samples from probenecid-treated healthy adults and identified 97 endogenous metabolites as putative OAT1/3 biomarkers (*34*). While these studies provide clinical evidence of identified biomarkers of OAT1/3, they are usually followed by separate *in vitro* assessments to mechanistically confirm the putative biomarkers as transporter substrates. The current study integrates all these components of biomarker discovery into one holistic approach to identify and qualify biomarkers of OAT1/3. We identified and qualified 23 features as robust biomarkers of OAT1/3 that varied in size, with *m/z* values ranging from 160 to 427 and potentially belong to different biochemical categories, including tryptophan metabolites, tyrosine metabolites, steroids, fatty acids, glucuronide conjugates, and sesquiterpenoids. Compared to baseline, the AUC of a feature with *m/z* 162.0550 was approximately 2.2-fold higher, and the CL_R_ was 60% lower, in the presence of probenecid. This biomarker was later characterized as I3CAG, which belongs to the tryptophan metabolic pathway. Similarly, kynurenic acid (*m/z* 190.0499) was validated as an OAT1/3 biomarker, with average AUC and CL_R_ ratios of 1.9 and 0.5, respectively. Biomarkers with *m/z* values of 204.0302, 283.0823, and 291.2319 potentially belong to the tryptophan metabolism, tyrosine metabolism and steroid pathways, respectively.

The known biomarkers of OAT1/3 are confounded by variability, selectivity, and sensitivity issues (*14, 16, 19*). For example, pyridoxic acid plasma concentrations are influenced by race, oral contraceptive usage, hypophosphatasia, and liver disease (*45–47*). 6β-HC is formed from cortisol via cytochrome P450 3A4 (CYP3A4) and can be affected by the variability in the enzyme activity (*48*). Similarly, GCDCA-S is a substrate of OATP1B1, complicating the renal *versus* liver DDI scenario (*16, 19*). An ideal biomarker should i) be selective for the transporter of interest; ii) not be affected by food, circadian rhythm or disease state; and iii) exhibit low inter and intraindividual variability, with consistent plasma concentrations across different days (*1, 23, 49*). The biomarker should be sensitive to transient inhibition of the pathway of interest, returning to baseline concentrations as inhibition diminishes. Metabolites with short half-lives have higher potential to be sensitive to transient inhibition and can be useful for predicting DDIs. While biomarkers with longer half-lives fail to show the effect of transient inhibition, these biomarkers can potentially be used to predict interindividual variability. The variability in the baseline plasma concentration of I3CAG was <30%. Plasma concentrations of this biomarker measured on two different days correlated significantly (data not shown), suggesting low interday variability in the baseline concentrations. Administration of probenecid led to a transient increase in biomarker concentrations, which decreased to the baseline levels with the elimination of probenecid. The precursor of I3CAG is a metabolite of the tryptophan pathway and is likely formed by the gut microbiota. Because the formation of glucuronide metabolite of I3CA could be influenced by the contribution of UGT enzymes, we identified UGT isoforms involved in I3CAG formation using recombinant enzyme systems. I3CA was determined to be a substrate of UGT1A6, UGT1A7 and UGT1A9 with comparable contributions from each isoform, suggesting that synthesis of the biomarker would not be limited by changes in the function of one or more UGTs. However, I3CAG is a substrate of OAT2, reducing the selectivity of the biomarker for OAT1/3. Given that the abundance of OAT2 is low in the kidneys (*41*), the overall contribution of OAT2 to renal elimination of I3CAG is minimal. OAT2 is abundant in liver (*42*), but the increase in plasma concentrations of I3CAG with a corresponding decrease in CL_R_ suggests that hepatic OAT2 is minimally affected by probenecid treatment. Further, strong and significant negative correlation of I3CAG pre-dose plasma concentration and furosemide plasma concentration fold change highlights the translational utility of this biomarker to prospectively predict OAT1/3-mediated DDI risk of substrate drugs. These data posit I3CAG as a sensitive, and selective OAT1/3 biomarker with minimal variability and baseline fluctuations.

The OAT1/3 biomarkers discovered in this study can potentially be used to investigate the OAT1/3 inhibitory potential of NCEs, wherein the biomarker concentrations can be measured during Phase 1 SAD and/or MAD studies, potentially eliminating the need for standalone clinical DDI studies using exogenous probe substrates (*49*). Once validated across different populations, these biomarkers could be used to assess changes in OAT1/3 activity in specific populations, including pediatric or geriatric groups, pregnant people, and patients with renal dysfunction and other disease conditions. However, the utility of a single biomarker in specific populations may be limited due to variability in the synthesis and/or elimination caused by factors such as age, pregnancy, and disease states (*23, 49*). Similarly, the single biomarker approach for predicting transporter-mediated DDIs during drug development is affected by the potential impact of the inhibitor on biomarker synthesis. To mitigate this limitation, a panel of biomarkers (e.g., 23 biomarkers of OAT1/3 identified in this study) representing different metabolic pathways would serve as a more robust predictor of interindividual variability and transporter inhibition. Furthermore, the MDBD approach can be extended to discover multiple biomarkers of other DMET proteins, which can then be utilized in a multiplexed panel to study the effects of NCEs on various drug absorption and disposition pathways.

Overall kidney function is determined by three processes: glomerular filtration, active secretion and reabsorption (*50*). Serum creatinine concentrations are commonly used as markers of glomerular filtration and clinically used as a surrogate for the overall kidney function (*51*). However, there is increasing evidence that as kidney function decreases, the effect on active secretion is not proportional with glomerular filtration (*52–55*). This raises questions about the utility of serum creatinine levels for determining kidney function, especially in cases of severe chronic kidney disease (*52*). The biomarkers of OAT1/3 discovered in this study, along with existing/new OAT1/3 and OCT2 biomarkers and creatinine, can potentially be used in a multiplex panel as non-invasive indicators of overall kidney function and process-specific functions.

In this study, high-resolution *m/z* values were used to predict metabolite identities. Further investigation is needed to characterize the structures of these biomarkers and determine the metabolic pathways involved in their synthesis. Once characterized, these biomarkers can be further assessed as substrates of OAT1/3 using *in vitro* transporter inhibition assays. Cross-validation of the current findings through multi-centric clinical studies using probenecid or other OAT1/3 inhibitors can help elevate the status of these biomarkers to *Tier 1* category as defined by the International Transporter Consortium (*1*). This study was conducted using healthy adults, opening the possibility to explore the effects of diet, age, sex, race, and other factors on the synthesis and elimination of these biomarkers. Because various extrinsic and intrinsic factors may influence the synthesis of these biomarkers, determining CL_R_ becomes key for evaluating the inhibition or differences in OAT1/3 activity. The confounding effect of MRP2/4-mediated apical efflux and shift in the rate-limiting step due to variable MRP2/4 function requires further investigation. The biomarker panel approach, once validated across specific populations, including pediatric or geriatric groups, pregnant people, patients with renal dysfunction or other diseases, can be used in clinic to assess changes in OAT1/3 function with translational utility to inform drug dosing and DDI prediction. Overall, the discovery of these OAT1/3 biomarkers opens new avenues for understanding the physiological and pathophysiological roles of these transporters, predicting DDIs during drug development, clinical monitoring of kidney function, and enabling precision dosing of OAT1/3 substrates in clinic.

## MATERIALS AND METHODS

### Materials

Indole-3-carboxylic acid (I3CA), bovine serum albumin (BSA), fetal bovine serum (FBS), and geneticin were obtained from Thermo Fisher Scientific (Rockford, IL). HPLC-grade methanol, MS-grade acetonitrile, formic acid, and water were procured from Fisher Chemical (Fair Lawn, NJ). Ammonium formate, indole-2-carboxylic acid (I2CA), alamethicin, furosemide, and UDP-glucuronic acid (UDPGA) were purchased from Sigma-Aldrich (St. Louis, MO). Probenecid, and furosemide-d5 were procured from Toronto Research Chemicals (Toronto, Canada). Testosterone glucuronide-d3 was obtained from Cerilliant Corporation (Round Rock, TX). Blasticidin and puromycin were obtained from InvivoGen (San Diego, CA). DMEM (high glucose, GlutaMAX^TM^ Supplement, pyruvate), Hank’s Balanced Salt Solution (HBSS), penicillin (10,000 Units/mL)/streptomycin (10,000 µg/mL) mixture, and phosphate buffered saline (PBS) were procured from Gibco (Amarillo, TX). HEPES was procured from Genesee Scientific (Morrisville, NC). L-proline and sodium butyrate were obtained from VWR (Radnor, PA). 10 mg/mL oral solution of furosemide and 500 mg tablets of probenecid were procured from West-Ward Pharmaceutical Corp. (Eatontown, NJ) and Lannett Company Inc. (Philadelphia, PA), respectively. Human embryonic kidney (HEK)-293 mock, OAT1, OAT2, and OAT3 over-expressing cells were provided by Charles River Laboratories (Budapest, Hungary). Wild-type Chinese hamster ovary (CHO) cells and CHO cells stably expressing human OATP1B1 or OATP1B3 were provided by Dr. Bruno Stieger (University of Zurich, Zurich, Switzerland). Recombinant human UGT (rUGT) enzymes were purchased from Corning Life Sciences (Riverfront, NY).

### Workflow of metabolomics-based DMET biomarker discovery (MDBD) approach

Probenecid inhibits renal OAT1/3, leading to an increase in plasma concentration and decrease in CL_R_ of OAT1/3 substrates. As per the MDBD approach (schematic shown in Fig. 1a), we conducted a clinical pharmacokinetic DDI study involving 16 healthy adult participants (8 males and 8 females) who were administered furosemide alone (baseline arm) or with probenecid (probenecid exposure arm) (Fig. 1b). Based on preliminary assessment of probenecid exposure, plasma samples (pool of 1, 1.5, and 2 h) and urine samples (0-4 h) representing maximum probenecid concentration (and thus maximum inhibitory potential) were selected for untargeted high resolution mass spectrometry (HRMS)-based metabolomics analysis. XCMS based data analysis yielded thousands of features (unique *m/z* values), which were analyzed using the paired t-test (baseline *versus* probenecid exposure arm) to identify biomarkers of OAT/3 (Fig. 1c). The identified biomarkers were mechanistically confirmed as substrates of OAT1/3 through *in vitro* transporter assays using transporter over-expressing cells. Biomarkers were screened as substrates of OAT1, OAT2, OAT3, OATP1B1, and OATP1B3. This was followed by clinical validation of the mechanistically confirmed biomarkers of OAT1/3, which included detailed 0-24 h pharmacokinetic analysis of plasma and urine samples from four study participants. Biomarkers with consistent pharmacokinetic profiles in the baseline and probenecid exposure arms were selected as clinically validated OAT1/3 biomarkers. Finally, the structure of one representative OAT1/3 biomarker was characterized using MS/MS fragmentation and further confirmed by spiking plasma with authentic reference standard.

### Clinical pharmacokinetic drug-drug interaction study

The Washington State University Institutional Review Board approved the clinical study protocol and informed consent form prior to subject recruitment. Study procedures were conducted at the Washington State University Human Research Clinic (HRC) on the Health Sciences Campus in accordance with the Code of Federal Regulations on the Protection of Human Subjects. The study was registered with the ClinicalTrials.gov database (NCT05365451). Subjects provided written informed consent prior to screening. Eligibility was determined based on medical history, routine clinical laboratory tests, physical examinations, pregnancy test for females of childbearing potential, and inclusion/exclusion criteria (Supplementary Table 1). Eight males and eight non-pregnant, non-lactating females (aged 26-64 years; detailed demographic information is provided in Supplementary Table 2) were enrolled in this open-label, 2-arm, crossover, fixed sequence pharmacokinetic study involving the OAT1/3 substrate furosemide and inhibitor probenecid (Fig. 1b). Considering the variability in furosemide pharmacokinetics and anticipated differences in pharmacokinetic measures (≥25%), a sample size of n=16 was optimum to achieve a power of 0.80 and an alpha value of 0.05. After an overnight fast, participants presented to the HRC in the morning of the inpatient study day, when their vital signs (heart rate, blood pressure, oxygen saturation) were recorded. Following placement of an indwelling venous catheter into an antecubital vein, they were administered a single oral dose (5 mg) of furosemide alone (baseline) or a single oral dose of probenecid (1,000 mg) followed 1 h later with 5 mg furosemide (probenecid exposure). They continued to fast during the next 4 h, when lunch was provided. Serial blood samples (5 mL) were collected before and 0.33, 0.6, 1, 1.5, 2, 2.5, 3, 3.5, 4, 5, 6, 8, 10, and 12 h after furosemide administration (Fig. 1b). Urine was collected into one or more jugs during the 0-4, 4-8, and 8-12 h intervals after furosemide administration. After the last blood draw, participants were discharged and provided jugs to collect their urine from 12-24 h. They returned to the HRC the next morning for a single blood draw at 24 h and to return their urine jug(s). Blood samples were cooled on ice prior to centrifugation to harvest plasma. Urine volume was measured using a graduated cylinder. Plasma and urine samples were stored at -80 and -20°C, respectively, until analysis.

### Quantification of furosemide and probenecid in plasma and urine samples

Plasma and urine samples as well as calibration standards were processed using an optimized protein precipitation approach as described in supplementary methods. The processed samples were analyzed using an M-class Waters ultra-performance liquid chromatography system coupled with Waters Xevo TQ-XS MS instrument using method details provided in supplementary methods, where the optimized multiple reaction monitoring parameters are listed in Supplementary Table 3.

### Pharmacokinetic and statistical analysis

LC-MS/MS data were analyzed using Skyline 23.1 (MacCoss Lab, Department of Genome Sciences, University of Washington, Seattle, WA). Urine volume was used to convert furosemide urine concentration into the amount excreted unchanged in urine. CL_R_ was calculated using equation 1.

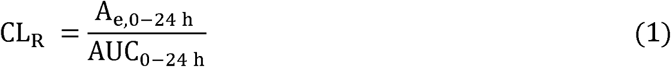

Statistical analysis was conducted using GraphPad Prism (v8.4.3; San Diego, CA). Plasma concentrations at individual time-points, AUC_0-24 h_, A_e,0-24 h_ and CL_R_ were compared using the paired t-test.

### Untargeted LC-MS metabolomics analysis of pooled plasma and urine samples

Pooled plasma was prepared by pooling 20-µL aliquots of plasma from three different time-points (1, 1.5, and 2 h). These time-points were based on the time of maximum probenecid concentration, leading to maximum inhibition of OAT1/3. The pooled plasma and urine (0-4 h) samples were processed as per the protocol described in Supplementary methods.

Untargeted metabolomics was performed using a Thermo EASY-nLC 1200 series system equipped with a Q-Exactive-HF MS instrument (San Jose, CA). The instrument operation details are provided in Supplementary methods.

### Biomarker identification: Untargeted metabolomics and statistical analysis

The MS raw files were converted to mzXML format using Raw Converter 1.2.0.1 software and the untargeted metabolomics data were analyzed using XCMS Online (Scripps Research, La Jolla, CA). Data analysis was performed using the paired t-test, with the baseline arm serving as the control and the probenecid exposure arm as the treatment groups. MS1 peak intensities were used as surrogates for plasma and urine concentrations, and detected features (*m/z* values) were filtered using biomarker identification criteria (Fig. 1c). Urine peak intensities were multiplied by urine volumes to estimate urine amounts. Putative biomarkers were screened based on a 1.5- to 10-fold increase in concentration in the presence of probenecid (p-value < 0.05). Features with more than a 10-fold increase in pooled plasma concentration were excluded as renal transporter-mediated DDIs are not expected to cause changes greater than this threshold (*56*). The apparent CL_R_ ratio was calculated using the following equations (2–4).

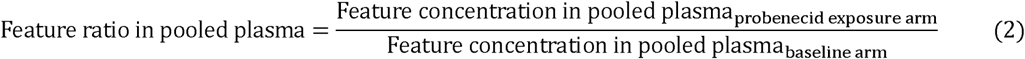

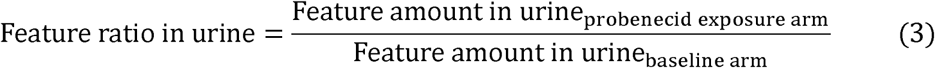

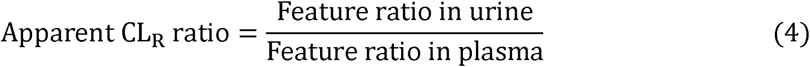

### Preparation of urine samples for *in vitro* transporter assay and *in vitro* transporter uptake assay in transporter-transfected cells

Urine samples from the baseline arm of the clinical study were pooled for use in transporter uptake assays. The urine samples underwent double protein precipitation (details described in Supplementary methods). Details pertaining to transporter uptake assays in HEK-293 mock, and OAT1, OAT2, or OAT3 over-expressing cells and CHO Wild-type and OATP1B1 or OATP1B3 over-expressing cells are provided in Supplementary methods.

### Untargeted metabolomics analysis of uptake assay data and comparison with list of identified OAT1/3 biomarkers

Data analysis of uptake assay samples was conducted using the unpaired t-test with mock cells representing the control and transporter over-expressing cells representing the treatment groups. Features with >1.5-fold higher uptake in OAT1 or OAT3 over-expressing cells (p-value < 0.05) were compared with the list of putative biomarkers identified from the clinical study samples. Features common to both the lists were confirmed as substrates of OAT1 and/or OAT3. For these features, the confounding effect of other transporters was assessed by determining their uptake in OAT2, OATP1B1, and/or OATP1B3 over-expressing cells.

### Clinical pharmacokinetic validation of putative biomarkers of OAT1/3

Of the 16 participants, 4 were selected for 0-24 h pharmacokinetic analysis based on maximum (n=2) and minimum (n=2) changes in furosemide pooled plasma concentration in the presence of probenecid. These samples were processed and analyzed similarly to pooled plasma and urine (0-4h) samples. MS1 peak intensities were used to depict concentrations, and peak intensities in urine samples were converted to amounts using urine volumes. The AUC and CL_R_ ratios were calculated using the following equations (5–6).

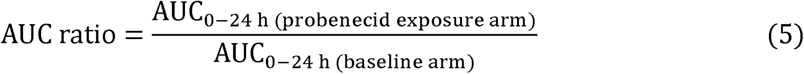

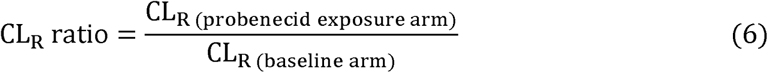

### MS parameters for high-resolution MS/MS analysis of analytes

MS/MS fragmentation analysis for structure characterization was carried out using Thermo Scientific EASY-nLC 1200 series system coupled with Q-Exactive-HF MS instrument. The MS was operated in full scan mode with *m/*z scan range of 100-675, resolution of 120,000, maximum injection time of 200 msec, and automatic gain control (AGC) target of 3×10^6^. For MS^2^, resolution of 30,000, maximum injection time of 100 msec, AGC target of 2×10^5^ and isolation window of 2 *m/z*. Stepped collision energy of 15, 20, and 30 eV was used for MS/MS fragmentation.

### Structure characterization of the feature with *m/*z 162.0550

For the case example feature with *m/z* 162.0550, the LC retention times and MS/MS fragmentation patterns of the biomarker and I2CA and I3CA standards were compared. The formation of ICAG from ICA by incubating the standards in human liver microsomes and recombinant human UGT enzymes was performed using protocol discussed in Supplementary methods (*57*). To distinguish if the glucuronide moiety in I3CAG is conjugated at the acyl or N-group, hydroxylamine derivatization assay was performed (*37*), as described in Supplementary methods.

### Biomarker panel approach

The top six biomarkers were chosen based on specific criteria: < 50% intraindividual variability and < 30% variability in baseline plasma concentrations (n=4 data points), lack of diurnal variations, ≥ 2-fold AUC ratio and ≥ 40% decrease in CL_R_. The fold changes in pooled plasma concentrations of these biomarkers in the 16 clinical study participants were averaged and correlated with the fold change in furosemide pooled plasma concentrations in the presence of probenecid.

## Supporting information

Supplementary Datafiles

## SUPPLEMENTARY MATERIAL

## SUPPLEMENTARY MATERIALS AND METHODS

**Preparation of plasma and urine samples for targeted analysis of furosemide and probenecid**

**Targeted LC-MS/MS analysis of furosemide and probenecid in plasma and urine samples**

**Sample preparation for untargeted metabolomics**

**Untargeted LC-MS/MS analysis of plasma and urine samples**

**Preparation of urine samples for *in vitro* transporter assay**

***In vitro* transporter uptake in transporter-transfected cells**

**Formation of indole-carboxylic acid glucuronide from indole-carboxylic acid**

**Hydroxylamine derivatization of indole-3-carboxylic acid glucuronide**

## SUPPLEMENTARY DATAFILES

**Supplementary Datafile 1.** List of *m/z* features significantly (a) increased (>1.5-fold) and (b) decreased (>1.5-fold) in pooled plasma samples on probenecid exposure.

**Supplementary Datafile 2.** List of *m/z* features with significantly higher uptake (>1.5-fold) in OAT1 over-expressing cells, compared to mock cells as detected in positive and negative ionization modes.

**Supplementary Datafile 3.** List of *m/z* features with significantly higher uptake (>1.5-fold) in OAT3 over-expressing cells, compared to mock cells as detected in positive and negative ionization modes.

**Supplementary Datafile 4.** *m/z* values of 57 features confirmed as substrates of either OAT1, OAT3, or both.

**Supplementary Datafile 5.** Pooled plasma fold change, apparent CL_R_ ratio, and *in vitro* transporter uptake (fold-change) data of validated OAT1/3 biomarkers.

**Supplementary Datafile 6.** Individual fold change (± standard deviation) values in pharmacokinetic endpoints of validated OAT1/3 biomarkers and furosemide on probenecid-mediated inhibition of renal OATs in healthy adult participants (n=4).

## SUPPLEMENTARY TABLES

**Supplementary Table 1.** The pharmacokinetic clinical probenecid-furosemide study inclusion and exclusion criteria.

**Supplementary Table 2.** Demographic characteristics of clinical study participants (n=16).

**Supplementary Table 3.** Optimized multiple reaction monitoring parameters used for the quantification of analytes.

## SUPPLEMENTARY FIGURES

**Supplementary Figure 1. Effect of probenecid treatment on plasma metabolome.**

**Supplementary Figure 2. Comparison of *m/z* features detected in pooled plasma (1, 1.5, 2 h) in negative ionization mode in the baseline and probenecid exposure arms.**

**Supplementary Figure 3. Comparison of *m/z* features detected in urine (0-4 h) in (a) positive and (b) negative ionization modes in the baseline and probenecid exposure arms.**

**Supplementary Figure 4. Comparison of *m/z* features concentrated in mock and OAT1 or OAT3 over-expressing cells following *in vitro* uptake assays in negative ionization mode.**

**Supplementary Figure 5. Uptake of validated OAT1/3 biomarkers and furosemide in OAT1 over-expressing cells compared to mock cells.**

**Supplementary Figure 6. Uptake of validated OAT1/3 biomarkers and furosemide in OAT3 over-expressing cells compared to mock cells.**

**Supplementary Figure 7. Fold change in pooled plasma concentration and apparent renal clearance (CL_R_) of validated OAT1/3 biomarkers and furosemide in the presence of probenecid.**

**Supplementary Figure 8. Pharmacokinetic profile of probenecid.**

**Supplementary Figure 9. Derivatization of indole-3-carboxylic acid glucuronide (I3CAG) to indole-3-hydroxamic acid.**

**Supplementary Figure 10. UGT isoforms involved in the formation of indole-3-carboxylic acid glucuronide (I3CAG).**

**Supplementary Figure 11. Correlation of indole-3-carboxylic acid pre-dose (0 min) plasma levels with furosemide pooled plasma fold change in healthy adult participants (n=16).**

**Supplementary Figure 12. Utility of a panel of top six validated OAT1/3 biomarkers in predicting interindividual variability in probenecid-mediated changes in furosemide plasma concentrations.**

## ACKNOWLEDGEMENTS

The authors thank Ms. Deena Hadi, Ms. Maxey Cherel, Dr. Vixen Cope, and Mr. Ryan Davy for their expert assistance in the completion of the clinical study. The authors thank Dr. Peter Turnbaugh for his insights and suggestions on the OAT1/3 biomarker, I3CAG.

## AUTHOR CONTRIBUTIONS

Designed the research: A.T., M.F.P., and B.P.

Conducted the clinical study: M.F.P.

Carried out the experiments: A.T. and K.D.H.

Carried out sample analysis: A.T. and D.K.S.

Performed data analysis: A.T. and B.P.

Contributed reagents/analytical tools: E.K., Z.G., T.T.D., and J.D.C

Wrote and reviewed the manuscript: A.T., D.K.S., K.D.H., Z.G., T.T.D., J.D.C., M.F.P. and B.P.

## REFERENCES

1. A. Galetin, K. L. R. Brouwer, D. Tweedie, K. Yoshida, N. Sjöstedt, L. Aleksunes, X. Chu, R. Evers, M. J. Hafey, Y. Lai, P. Matsson, A. Riselli, H. Shen, A. Sparreboom, M. V. S. Varma, J. Yang, X. Yang, S. W. Yee, M. J. Zamek-Gliszczynski, L. Zhang, K. M. Giacomini, Membrane transporters in drug development and as determinants of precision medicine. Nat Rev Drug Discov 23, 255–280 (2024).

2. S.-C. Lee, V. Arya, X. Yang, D. A. Volpe, L. Zhang, Evaluation of transporters in drug development: Current status and contemporary issues. Adv Drug Deliv Rev 116, 100–118 (2017).

3. ICH M12 Guideline: Drug Interaction Studies. https://www.fda.gov/regulatory-information/search-fda-guidance-documents/m12-drug-interaction-studies.

4. FDA Guidance for Industry: In Vitro Drug Interaction Studies — Cytochrome P450 Enzyme- and Transporter-Mediated Drug Interactions. https://www.fda.gov/media/134582/download.

5. European Medicines Agency: Guideline on the investigation of drug interactions. https://www.ema.europa.eu/en/investigation-drug-interactions-scientific-guideline.

6. A. D. Rodrigues, Reimagining the framework supporting the static analysis of transporter drug interaction risk; integrated use of biomarkers to generate pan□transporter inhibition signatures. Clin Pharmacol Ther (2022), doi:10.1002/cpt.2713.

7. R. Kikuchi, P. P. Chothe, X. Chu, F. Huth, K. Ishida, N. Ishiguro, R. Jiang, H. Shen, S. H. Stahl, M. V. S. Varma, M. Willemin, B. L. Morse, Utilization of OATP1B Biomarker Coproporphyrin□I to Guide Drug–Drug Interaction Risk Assessment: Evaluation by the Pharmaceutical Industry. Clin Pharmacol Ther 114, 1170–1183 (2023).

8. H. V. Kalluri, R. Kikuchi, S. Coppola, J. Schmidt, M. F. Mohamed, D. A. J. Bow, A. H. Salem, Coproporphyrin I Can Serve as an Endogenous Biomarker for OATP1B1 Inhibition: Assessment Using a Glecaprevir/Pibrentasvir Clinical Study. Clin Transl Sci 14, 373–381 (2021).

9. Y. Lai, The Role of Coproporphyrins As Endogenous Biomarkers for Organic Anion Transporting Polypeptide 1B Inhibition–Progress from 2016 to 2023. Drug Metabolism and Disposition 51, 950–961 (2023).

10. S. K. Nigam, W. Wu, K. T. Bush, M. P. Hoenig, R. C. Blantz, V. Bhatnagar, Handling of drugs, metabolites, and uremic toxins by kidney proximal tubule drug transporters. Clin J Am Soc Nephrol 10, 2039–2049 (2015).

11. T. Sekine, S. H. Cha, H. Endou, The multispecific organic anion transporter (OAT) family. Pflugers Arch 440, 337–350 (2000).

12. G. Burckhardt, Drug transport by Organic Anion Transporters (OATs)Pharmacol Ther 136, 106–130 (2012).

13. J. Tang, H. Shen, X. Zhao, V. K. Holenarsipur, T. T. Mariappan, Y. Zhang, E. Panfen, J. Zheng, W. G. Humphreys, Y. Lai, Endogenous plasma kynurenic acid in human: A newly discovered biomarker for drug-drug interactions involving organic anion transporter 1 and 3 inhibition. Drug Metab Dispos 49, 1063–1069 (2021).

14. H. Shen, V. K. Holenarsipur, T. T. Mariappan, D. M. Drexler, J. L. Cantone, P. Rajanna, S. S. Gautam, Y. Zhang, J. Gan, P. A. Shipkova, P. Marathe, W. G. Humphreys, Evidence for the validity of pyridoxic acid (PDA) as a plasma-based endogenous probe for OAT1 and OAT3 function in healthy subjects. J Pharmacol Exp Ther 368, 136–145 (2019).

15. H. Shen, D. M. Nelson, R. V. Oliveira, Y. Zhang, C. A. McNaney, X. Gu, W. Chen, C. Su, M. D. Reily, P. A. Shipkova, J. Gan, Y. Lai, P. Marathe, W. G. Humphreys, Discovery and validation of pyridoxic acid and homovanillic acid as novel endogenous plasma biomarkers of organic anion transporter (OAT) 1 and OAT3 in Cynomolgus monkeys. Drug Metab Dispos 46, 178–188 (2018).

16. M. E. Willemin, T. K. Van Der Made, I. Pijpers, L. Dillen, A. Kunze, S. Jonkers, K. Steemans, A. Tuytelaars, F. Jacobs, M. Monshouwer, D. Scotcher, A. Rostami-Hodjegan, A. Galetin, J. Snoeys, Clinical investigation on endogenous biomarkers to predict strong OAT-mediated drug-drug interactions. Clin Pharmacokinet 60, 1187–1199 (2021).

17. Y. Tsuruya, K. Kato, Y. Sano, Y. Imamura, K. Maeda, Y. Kumagai, Y. Sugiyama, H. Kusuhara, Investigation of endogenous compounds applicable to drug-drug interaction studies involving the renal organic anion transporters, OAT1 and OAT3, in humans. Drug Metab Dispos 44, 1825–1933 (2016).

18. Y. Imamura, Y. Tsuruya, K. Damme, D. Heer, Y. Kumagai, K. Maeda, N. Murayama, N. Okudaira, A. Kurihara, T. Izumi, Y. Sugiyama, H. Kusuhara, 6β-Hydroxycortisol Is an endogenous probe for evaluation of drug-drug interactions involving a multispecific renal organic anion transporter, OAT3/SLC22A8, in healthy subjects. Drug Metab Dispos 42, 685–694 (2014).

19. I. Takehara, H. Terashima, T. Nakayama, T. Yoshikado, M. Yoshida, K. Furihata, N. Watanabe, K. Maeda, O. Ando, Y. Sugiyama, H. Kusuhara, Investigation of glycochenodeoxycholate sulfate and chenodeoxycholate glucuronide as surrogate endogenous probes for drug interaction studies of OATP1B1 and OATP1B3 in healthy Japanese volunteers. Pharm Res 34, 1601–1614 (2017).

20. E. P. Rhee, R. E. Gerszten, Metabolomics and Cardiovascular Biomarker Discovery. Clin Chem 58, 139–147 (2012).

21. W. J. Griffiths, T. Koal, Y. Wang, M. Kohl, D. P. Enot, H. Deigner, Targeted Metabolomics for Biomarker Discovery. Angewandte Chemie International Edition 49, 5426–5445 (2010).

22. W. R. Wikoff, M. A. Nagle, V. L. Kouznetsova, I. F. Tsigelny, S. K. Nigam, Untargeted Metabolomics Identifies Enterobiome Metabolites and Putative Uremic Toxins as Substrates of Organic Anion Transporter 1 (Oat1). J Proteome Res 10, 2842–2851 (2011).

23. X. Chu, M. Liao, H. Shen, K. Yoshida, A. A. Zur, V. Arya, A. Galetin, K. M. Giacomini, I. Hanna, H. Kusuhara, Y. Lai, D. Rodrigues, Y. Sugiyama, M. J. Zamek-Gliszczynski, L. Zhang, Clinical probes and endogenous biomarkers as substrates for transporter drug-drug interaction evaluation: Perspectives from the international transporter consortium. Clin Pharmacol Ther 104, 836–864 (2018).

24. S. A. Eraly, V. Vallon, D. A. Vaughn, J. A. Gangoiti, K. Richter, M. Nagle, J. C. Monte, T. Rieg, D. M. Truong, J. M. Long, B. A. Barshop, G. Kaler, S. K. Nigam, Decreased renal organic anion secretion and plasma accumulation of endogenous organic anions in OAT1 knock-out mice. J Biol Chem 281, 5072–5083 (2006).

25. X. Gou, F. Ran, J. Yang, Y. Ma, X. Wu, Construction and Evaluation of a Novel Organic Anion Transporter 1/3 CRISPR/Cas9 Double-Knockout Rat Model. Pharmaceutics 14 (2022), doi:10.3390/pharmaceutics14112307.

26. W. Wu, N. Jamshidi, S. A. Eraly, H. C. Liu, K. T. Bush, B. O. Palsson, S. K. Nigam, Multispecific drug transporter Slc22a8 (Oat3) regulates multiple metabolic and signaling pathways. Drug Metab Dispos 41, 1825–1834 (2013).

27. V. Vallon, S. A. Eraly, W. R. Wikoff, T. Rieg, G. Kaler, D. M. Truong, S.-Y. Ahn, N. R. Mahapatra, S. K. Mahata, J. A. Gangoiti, W. Wu, B. A. Barshop, G. Siuzdak, S. K. Nigam, Organic Anion Transporter 3 Contributes to the Regulation of Blood Pressure. Journal of the American Society of Nephrology 19, 1732–1740 (2008).

28. W. Wu, K. T. Bush, S. K. Nigam, Key Role for the Organic Anion Transporters, OAT1 and OAT3, in the in vivo Handling of Uremic Toxins and Solutes. Sci Rep 7 (2017), doi:10.1038/s41598-017-04949-2.

29. S. A. Eraly, V. Vallon, T. Rieg, J. A. Gangoiti, W. R. Wikoff, G. Siuzdak, B. A. Barshop, S. K. Nigam, Multiple organic anion transporters contribute to net renal excretion of uric acid. Physiol Genomics 33, 180–192 (2008).

30. J. C. Granados, V. Ermakov, K. Maity, D. R. Vera, G. Chang, S. K. Nigam, The kidney drug transporter OAT1 regulates gut microbiome–dependent host metabolism. JCI Insight 8 (2023), doi:10.1172/jci.insight.160437.

31. Y. Ma, F. Ran, M. Xin, X. Gou, X. Wang, X. Wu, Albumin-bound kynurenic acid is an appropriate endogenous biomarker for assessment of the renal tubular OATs-MRP4 channel. J Pharm Anal (2023), doi:10.1016/j.jpha.2023.05.007.

32. A. Thakur, V. Saradhi Mettu, D. K. Singh, B. Prasad, Effect of probenecid on blood levels and renal elimination of furosemide and endogenous compounds in rats: Discovery of putative organic anion transporter biomarkers. Biochem Pharmacol 218, 115867 (2023).

33. S. Yee, M. Giacomini, C. Hsueh, D. Weitz, X. Liang, S. Goswami, J. Kinchen, A. Coelho, A. Zur, K. Mertsch, W. Brian, D. Kroetz, K. Giacomini, Metabolomic and Genome□wide Association Studies Reveal Potential Endogenous Biomarkers for OATP1B1. Clin Pharmacol Ther 100, 524–536 (2016).

34. J. C. Granados, V. Bhatnagar, S. K. Nigam, Blockade of organic anion transport in humans after treatment with the drug probenecid leads to major metabolic alterations in plasma and urine. Clin Pharmacol Ther 112, 653–664 (2022).

35. A. Selen, G. L. Amidon, P. G. Wellingx, Pharmacokinetics of Probenecid Following Oral Doses to Human Volunteers. J Pharm Sci 71, 1238–1242 (1982).

36. HMDB website. https://hmdb.ca/metabolites/HMDB0013189.

37. A. D. N. Vaz, W. W. Wang, A. J. Bessire, R. Sharma, A. E. Hagen, A rapid and specific derivatization procedure to identify acyl□glucuronides by mass spectrometry. Rapid Communications in Mass Spectrometry 24, 2109–2121 (2010).

38. V. Tolstikov, A. J. Moser, R. Sarangarajan, N. R. Narain, M. A. Kiebish, Current Status of Metabolomic Biomarker Discovery: Impact of Study Design and Demographic Characteristics. Metabolites 10, 224 (2020).

39. J. Xia, D. I. Broadhurst, M. Wilson, D. S. Wishart, Translational biomarker discovery in clinical metabolomics: an introductory tutorial. Metabolomics 9, 280–299 (2013).

40. F. A. Castelli, G. Rosati, C. Moguet, C. Fuentes, J. Marrugo-Ramírez, T. Lefebvre, H. Volland, A. Merkoçi, S. Simon, F. Fenaille, C. Junot, Metabolomics for personalized medicine: the input of analytical chemistry from biomarker discovery to point-of-care tests. Anal Bioanal Chem 414, 759–789 (2022).

41. A. Thakur, G. Yue, D. Ahire, V. S. Mettu, A. Al Maghribi, K. Ford, L. Peixoto, J. S. Leeder, B. Prasad, Sex and the Kidney Drug□Metabolizing Enzymes and Transporters: Are Preclinical Drug Disposition Data Translatable to Humans? Clin Pharmacol Ther 116, 235– 246 (2024).

42. A. Vildhede, E. Kimoto, A. D. Rodrigues, M. V. S. Varma, Quantification of Hepatic Organic Anion Transport Proteins OAT2 and OAT7 in Human Liver Tissue and Primary Hepatocytes. Mol Pharm 15, 3227–3235 (2018).

43. M. J. Zamek-Gliszczynski, K. M. Goldstein, A. Paulman, T. K. Baker, T. P. Ryan, Minor Compensatory Changes in SAGE Mdr1a (P-gp), Bcrp, and Mrp2 Knockout Rats Do Not Detract from Their Utility in the Study of Transporter-Mediated Pharmacokinetics. Drug Metab Dispos 41, 1174–1178 (2013).

44. A. Basit, Z. Radi, V. S. Vaidya, M. Karasu, B. Prasad, Kidney Cortical Transporter Expression across Species Using Quantitative Proteomics. Drug Metabolism and Disposition 47, 802–808 (2019).

45. S. P. Coburn, K. G. Thampy, H. W. Lane, P. S. Conn, P. J. Ziegler, D. L. Costill, J. D. Mahuren, W. J. Fink, D. R. Pearson, W. E. Schaltenbrand, Pyridoxic acid excretion during low vitamin B-6 intake, total fasting, and bed rest. Am J Clin Nutr 62, 979–983 (1995).

46. S. P. Coburn, R. D. Reynolds, J. D. Mahuren, W. E. Schaltenbrand, Y. Wang, K. L. Ericson, M. P. Whyte, Y. M. Zubovic, P. J. Ziegler, D. L. Costill, W. J. Fink, D. R. Pearson, T. A. Pauly, K. G. Thampy, J. Wortsman, Elevated plasma 4-pyridoxic acid in renal insufficiency. Am J Clin Nutr 75, 57–64 (2002).

47. E. A. Donald, T. R. Bossé, The vitamin B6 requirement in oral contraceptive users II. Assessment by tryptophan metabolites, vitamin B6, and pyridoxic acid levels in urine. Am J Clin Nutr 32, 1024–1032 (1979).

48. C.-C. Peng, I. Templeton, K. E. Thummel, C. Davis, K. L. Kunze, N. Isoherranen, Evaluation of 6β-Hydroxycortisol, 6β-Hydroxycortisone, and a Combination of the Two as Endogenous Probes for Inhibition of CYP3A4 In Vivo. Clin Pharmacol Ther 89, 888–895 (2011).

49. H. J. Choi, S. Madari, F. Huang, Utilising Endogenous Biomarkers in Drug Development to Streamline the Assessment of Drug–Drug Interactions Mediated by Renal Transporters: A Pharmaceutical Industry Perspective. Clin Pharmacokinet 63, 735–749 (2024).

50. S. E. Tett, C. M. J. Kirkpatrick, A. S. Gross, A. J. McLachlan, Principles and clinical application of assessing alterations in renal elimination pathways. Clin Pharmacokinet 42, 1193–1211 (2003).

51. K. Kashani, M. H. Rosner, M. Ostermann, Creatinine: From physiology to clinical application. Eur J Intern Med 72, 9–14 (2020).

52. S. P. F. Tan, D. Scotcher, A. Rostami□Hodjegan, A. Galetin, Effect of Chronic Kidney Disease on the Renal Secretion via Organic Anion Transporters 1/3: Implications for Physiologically□Based Pharmacokinetic Modeling and Dose Adjustment. Clin Pharmacol Ther 112, 643–652 (2022).

53. H. Takita, D. Scotcher, R. Chinnadurai, P. A. Kalra, A. Galetin, Physiologically□Based Pharmacokinetic Modelling of Creatinine□Drug Interactions in the Chronic Kidney Disease Population. CPT Pharmacometrics Syst Pharmacol 9, 695–706 (2020).

54. A. Chapron, D. Shen, B. Kestenbaum, C. Robinson□Cohen, J. Himmelfarb, C. Yeung, Does Secretory Clearance Follow Glomerular Filtration Rate in Chronic Kidney Diseases? Reconsidering the Intact Nephron Hypothesis. Clin Transl Sci 10, 395–403 (2017).

55. S. Pradhan, S. B. Duffull, R. J. Walker, D. F. B. Wright, The intact nephron hypothesis as a model for renal drug handling. Eur J Clin Pharmacol 75, 147–156 (2019).

56. B. Feng, M. V. Varma, Evaluation and Quantitative Prediction of Renal Transporter□Mediated Drug□Drug Interactions. The Journal of Clinical Pharmacology 56 (2016), doi:10.1002/jcph.702.

57. D. Ahire, S. Heyward, B. Prasad, Intestinal Metabolism of Diclofenac by Polymorphic UGT2B17 Correlates with its Highly Variable Pharmacokinetics and Safety across Populations. Clin Pharmacol Ther 114, 161–172 (2023).

