## Supplementary Information for "From discovery to translation: Endogenous substrates of OAT1 and OAT3 as clinical biomarkers for renal secretory function"

**SUPPLEMENTARY MATERIALS AND METHODS**

**Preparation of plasma and urine samples for targeted analysis of furosemide and probenecid**

To the 100 µL of plasma samples, 400 µL of ice-cold acetonitrile containing 100 ng/mL furosemide-d5 as internal standard was added, vortexed, and centrifuged (16,000 xg and 4 °C for 10 min). The supernatant was dried at 45 °C for 2 h, then reconstituted with 100 µL of acetonitrile:1 mM ammonium formate in water (30:70, v/v), vortexed, and centrifuged (16,000 xg and 4 °C for 10 min). The supernatant was transferred into an LC-MS vial for analysis of furosemide and probenecid. Calibration standards (10x concentrations) were prepared by serially diluting the stock solutions of furosemide, and probenecid using methanol:water (80:20, v/v). Calibration solutions were prepared using 2% BSA in PBS as matrix.

To the 50 µL of urine samples, 50 µL of ice-cold acetonitrile containing 4000 ng/mL furosemide-d5 as internal standard was added, vortexed, and centrifuged (16,000 xg and 4 °C for 10 min). Thereafter, 80 µL of supernatant was diluted with 320 µL of 1 mM ammonium formate in water, vortexed, followed by addition of 100 µL of acetonitrile, and centrifuged (16,000 xg and 4 °C for 10 min). The supernatant was transferred into an LC-MS vial for analysis of furosemide. Calibration standard solutions of furosemide were prepared in water.

**Targeted LC-MS/MS analysis of furosemide and probenecid in plasma and urine samples**

The processed samples and calibration standards were analyzed using an M-class Waters ultra-performance liquid chromatography system coupled with Waters Xevo TQ-XS MS instrument equipped with an electrospray ionization source. Chromatographic separation was achieved using Acquity UPLC^®^ HSS T3 (1.8 µm, 100x1 mm) column set at 40 ºC. The flow rate was 50 µL/min, with 0.1% formic acid in water and 0.1% formic acid in acetonitrile as mobile phases A and B, respectively. The mobile phase gradient program (%B) was: 0-1 min (5%), 1-2.5 min (5 to 55%), 2.5-4.2 min (55 to 68%), 4.2-5.2 min (68 to 90%), 5.2-8 (90%), 8-8.5 min (90 to 5%) and 8.5-11.5 min (5%) for sample analysis. The mass spectrometer was operated in the negative ionization mode for the quantification of furosemide and probenecid. Optimized multiple reaction monitoring parameters are provided in Supplementary Table 9.

**Sample preparation for untargeted metabolomics**

Pooled plasma was prepared by pooling 20-µL aliquots of plasma from three different time-points (1, 1.5, and 2 h). These time-points were based on the time of maximum probenecid concentration, leading to maximum inhibition of OAT1/3. To the 60 µL of pooled plasma and urine (0-4 h sample), 400 µL of ice-cold acetonitrile were added (first protein precipitation step), vortexed, and centrifuged (16,000 xg and 4 °C for 10 min). The supernatant was dried at 45 °C for 2 h, then reconstituted with 60 µL of water, vortexed, and subjected to a second protein precipitation step by addition of 400 µL of ice-cold acetonitrile, followed by overnight storage at -20 °C. The next day, the samples were thawed, vortexed, and centrifuged (16,000 xg and 4 °C for 10 min). The supernatant was dried at 45 °C for 2 h, and the pellet was reconstituted with 100 µL of acetonitrile:1 mM ammonium formate in water (10:90, v/v), vortexed, and centrifuged (16,000 xg and 4 °C for 10 min). The supernatant was transferred into an LC-MS vial for untargeted metabolomics analysis.

**Untargeted LC-MS/MS analysis of plasma and urine samples**

Untargeted metabolomics was performed using a Thermo EASY-nLC 1200 series system equipped with a Q-Exactive-HF MS instrument (San Jose, CA). LC separation was achieved using a Thermo Scientific Pep-Map^TM^ RSLC C18 (2 µm, 250 x 0.075 mm) column set at 40 °C with a mobile phase flow rate of 300 nL/min and an injection volume of 1 µL. Mobile phase A and B consisted of 0.1% formic acid in water and 0.1% formic acid in 80% acetonitrile, respectively, with a gradient program (%B) as follows: 0 to 1 min (10%), 1 to 11 min (10-60%), 11 to 20 min (60-80%), 20 to 25 min (80-100%), and 25 to 40 min (100%). MS ionization was conducted using an EasySpray ionization source, and sample analysis was performed in both positive and negative ionization polarities with data-independent acquisition mode. The MS was operated in full scan mode with *m/z* scan range of 70-750, resolution of 120,000, capillary temperature of 300 °C, S-lens RF level of 50, and maximum injection time of 150 ms.

**Preparation of urine samples for *in vitro* transporter assay**

Urine samples from the baseline arm of the clinical study were pooled for use in transporter uptake assays. The urine samples underwent double protein precipitation (as described previously) and were placed overnight in a freezer at -20 °C. The following day, the samples were thawed, vortexed, and centrifuged at 16,000 xg and 4 °C for 10 min. The supernatant was dried at 45 °C for 2 h and the pellet was reconstituted with different volumes of HBSS (<0.5% methanol) to create three types of samples: 5-fold diluted (0.2x), undiluted (1x), and 5-fold concentrated (5x). These samples were incubated with cells during the uptake assays.

***In vitro* transporter uptake in transporter-transfected cells**

HEK-293 mock cells and OAT1, OAT2 or OAT3 over-expressing cells were cultured in DMEM supplemented with 10% FBS, penicillin, and streptomycin. Blasticidin was included as a selection antibiotic for OAT1 over-expressing cells, while puromycin was used for OAT2 and OAT3 over-expressing cells. Cells were seeded onto 24-well poly D-lysine coated plates at a density of 500,000 cells per well and maintained for 24 h at 37 °C, 5% CO_2_ and 90% relative humidity. For the assay, the medium was aspirated, and the cells were washed twice with 300 µL of pre-warmed PBS. The cells were then acclimatized by incubating with pre-warmed PBS for 15 min at 37 °C, 5% CO_2_. Uptake was initiated by incubating the urine mix with the cells for 5 min, followed by washing the cells three times with ice-cold PBS to terminate cellular uptake. The cells were lysed by adding 600 µL of ice-cold acetonitrile and incubating at 4 °C for 20 min. The lysates were centrifuged at 16,000 xg and 4 °C for 10 min, dried at 45 °C, and the pellet was reconstituted with 50 µL of acetonitrile:1 mM ammonium formate in water (10:90, v/v) and then transferred into LC-MS vials for untargeted metabolomics analysis. Wild-type CHO cells and CHO cells stably expressing human OATP1B1 (passage 13) or OATP1B3 (passages 29) were maintained at 37 °C in a humidified 5% CO_2_ atmosphere in DMEM (low-glucose) containing 25 mM HEPES supplemented with 10% FBS and 50 mg/ml L-proline culture without geneticin (wild-type cells) or with 0.5 mg/ml geneticin (OATP-expressing cells). Cells were seeded onto 12-well plates at a density of 0.3 × 10^6^ cells per well. Transporter expression was induced by incubating cells with 10 mM sodium butyrate for 24 h prior to transport experiment. Waymouth transport buffer (135 mM sodium chloride, 1.3 mM HEPES, 2.8 mM D-glucose, 0.5 mM potassium chloride, 0.25 mM calcium dichloride, 0.12 mM magnesium dichloride, 80 mM magnesium sulfate, pH 7.4) was used in place of culture medium for all transport experiments. Prior to initiating transport, cells were washed three times with prewarmed transport buffer and allowed to equilibrate for 5 min at 37 °C. Transport was carried out for 5 min, then terminated using three ice cold transport buffer washes, then the cells were lysed with ice-cold acetonitrile.

Initially, 0.2x, 1x, and 5x urine were used for the uptake assay, and the uptake of positive controls (furosemide and kynurenic acid) was monitored. Based on these results, 0.2x and 1x urine samples were finally used. These concentrations were optimized based on the ability of substrates to inhibit the uptake of other substrates present in the urine.

**Formation of indole-carboxylic acid glucuronide from indole-carboxylic acid**

I2CA and I3CA standards (50 µM) were incubated with human liver microsomes (10 µg) in the presence of alamethicin (0.1 mg/mL), BSA (0.01%), and UDPGA (2.5 mM) at 37 °C for 1 h. The reaction was quenched by adding acetonitrile, vortex mixing for 5 min, centrifuging (16,000 xg and 4°C for 10 min), then transferring to LC-MS vials for high-resolution MS analysis (see MS parameters for high-resolution MS/MS analysis above). Similarly, 50 µM I3CA was incubated with 10 µg each of 13 recombinant UGT enzymes (UGT1A1, UGT1A3, UGT1A4, UGT1A6, UGT1A7, UGT1A8, UGT1A9, UGT1A10, UGT2B4, UGT2B7, UGT2B10, UGT2B15, and UGT2B17) and the formation of I3CAG was monitored in each system. The reaction was quenched by adding acetonitrile (containing 50 ng/mL testosterone glucuronide-d3 as an internal standard), vortex mixing for 5 min, centrifuging, then transferring to LC-MS vials for analysis. I3CA was used as a surrogate to measure I3CAG and optimized multiple reaction monitoring parameters are provided in Supplementary Table 9.

**Hydroxylamine derivatization of indole-3-carboxylic acid glucuronide**

Urine samples were processed as described above. To 100 µL of urine sample, an equal volume of 50% aqueous hydroxylamine solution was added, and reacted at room temperature for 10 h, and then directly injected for LC-MS analysis. The formation of indole-3-hydroxamic acid was determined by HRMS analysis using the parameter described above.

**SUPPLEMENTARY DATAFILES**

**Supplementary Datafile 1.** List of *m/z* features significantly (a) increased (>1.5-fold) and (b) decreased (>1.5-fold) in pooled plasma samples on probenecid exposure.

**Supplementary Datafile 2.** List of *m/z* features with significantly higher uptake (>1.5-fold) in OAT1 over-expressing cells, compared to mock cells as detected in positive and negative ionization modes.

**Supplementary Datafile 3.** List of *m/z* features with significantly higher uptake (>1.5-fold) in OAT3 over-expressing cells, compared to mock cells as detected in positive and negative ionization modes.

**Supplementary Datafile 4.** *m/z* values of 57 features confirmed as substrates of either OAT1, OAT3, or both.

**Supplementary Datafile 5.** Pooled plasma fold change, apparent CLR ratio, and in vitro transporter uptake (fold-change) data of validated OAT1/3 biomarkers.

**Supplementary Datafile 6.** Individual fold change (± standard deviation) values in pharmacokinetic endpoints of validated OAT1/3 biomarkers and furosemide on probenecid-mediated inhibition of renal OATs in healthy adult participants (n=4).

**SUPPLEMENTARY TABLES**

**Supplementary Table 1.** The pharmacokinetic clinical probenecid-furosemide study inclusion and exclusion criteria.

**Inclusion criteria:**

Aged from 18-65 years and healthy

Not taking any medications (prescription and non-prescription) or dietary/herbal supplements known to alter the pharmacokinetics of any study drug

Willing to abstain from consuming any alcoholic beverages for one day prior to the inpateint visiti and each of the outpatient visits

Willing to abstain from consuming caffeinated beverages or other caffeine-containing products the evening before the inpatient visit

Ability to understand the informed consent form

**Exclusion criteria:**

Children aged 17 years or less and adults aged 66 years or more

Any current major illness or chronic illness such as kidney disease, hepatic disease, diabetes mellitus, hypertension, coronary artery disease, chronic obstructive pulmonary diease, cancer, or HIV/AIDS

History of drug or alcohol addiction or major psychiatric illness

Pregnant or nursing females

History of intolerance or allergy to furosemide or probenecid

Taking concomitant medications, both prescription and non-prescription (including dietary supplements/herbal products) known to alter the pharmacokinetics of furosemide or probenecid

**Supplementary Table 2.** Demographic characteristics of clinical study participants (n=16).

| **Subject ID** | **Age (years)** | **Sex** | **Race^a^** | **Ethnicity^a^** |
| --- | --- | --- | --- | --- |
| 1 | 28 | Female | White | Not Hispanic or Latino |
| 2 | 58 | Female | White | Not Hispanic or Latino |
| 3 | 39 | Male | White | Unknown or not reported |
| 4 | 27 | Female | Unknown not reported | Unknown or not reported |
| 5 | 26 | Female | White | Not Hispanic or Latino |
| 6 | 36 | Male | White | Not Hispanic or Latino |
| 7 | 57 | Female | White | Unknown or not reported |
| 8 | 31 | Male | White | Not Hispanic or Latino |
| 9 | 27 | Male | Black or African American | Not Hispanic or Latino |
| 10 | 64 | Female | White | Not Hispanic or Latino |
| 11 | 56 | Male | White | Not Hispanic or Latino |
| 12 | 24 | Female | White | Not Hispanic or Latino |
| 13 | 28 | Female | Asian | Not Hispanic or Latino |
| 14 | 35 | Male | Asian | Not Hispanic or Latino |
| 15 | 30 | Male | More than one race | Not Hispanic or Latino |
| 16 | 50 | Male | White | Not Hispanic or Latino |
| ^a^Self-reported via a standard form | | |  |  |

**Supplementary Table 3.** Optimized multiple reaction monitoring parameters used for the quantification of analytes.

| **Compound** | **Polarity** | **Precursor ion (*m/z)*** | **Product ion (*m/z*)** | **Cone voltage (V)** | **Collision energy (eV)** |
| --- | --- | --- | --- | --- | --- |
| Furosemide | Negative | 329 | 205, 285 | 25 | 20, 15 |
| Furosemide-d5 | Negative | 334 | 290 | 25 | 20 |
| Probenecid | Negative | 284.1 | 140, 240 | 25 | 24, 15 |
| Indole-3-carboxylic acid | Positive | 162.1 | 118.1 | 25 | 13 |

**SUPPLEMENTARY FIGURES**

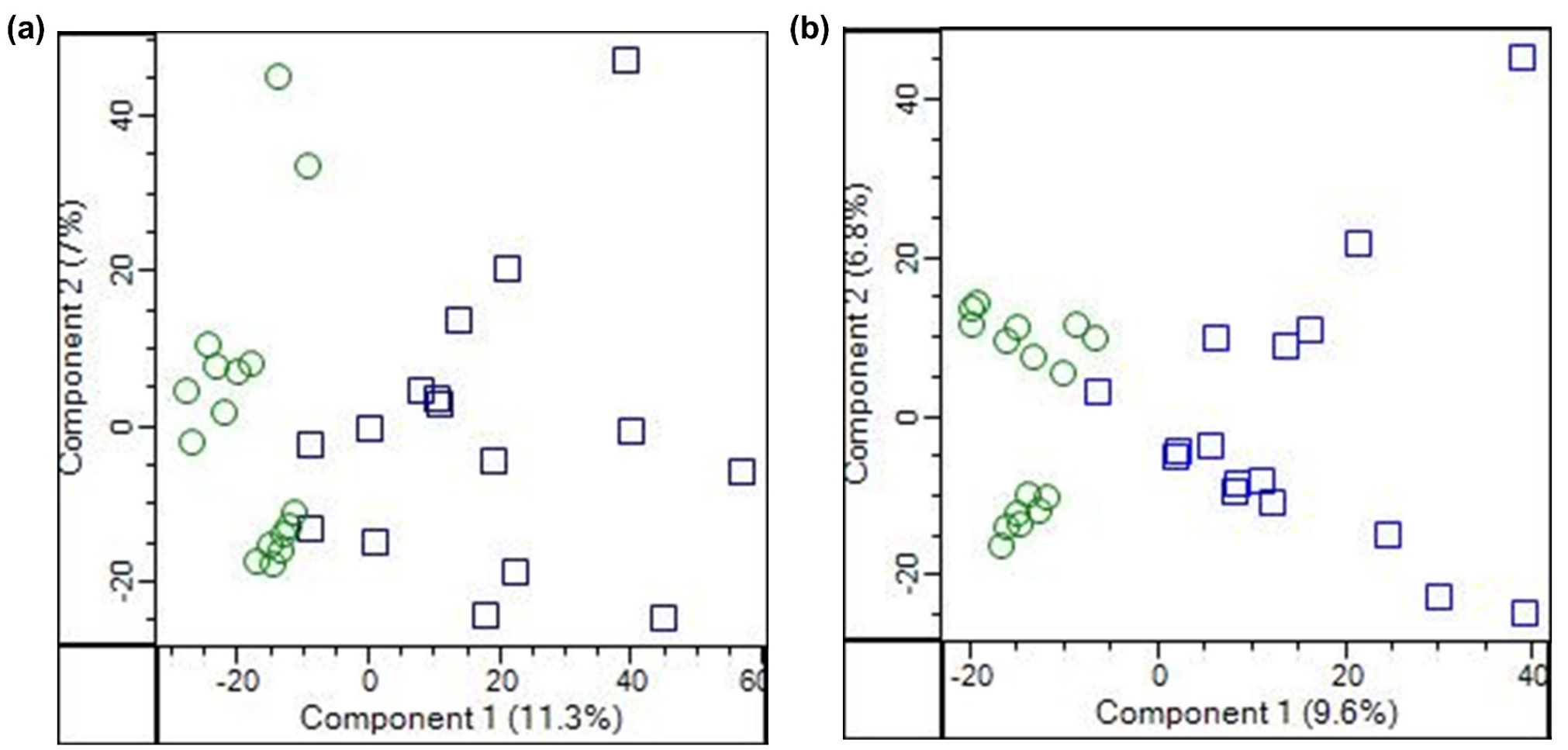

**Supplementary Figure 1. Effect of probenecid treatment on plasma metabolome.** Principal component analysis highlighting effects of probenecid treatment on the global plasma metabolome, as revealed in both (a) positive, and (b) negative ionization modes. Each green circle represents the global metabolome in a study participant in baseline (furosemide alone) arm with blue squares representing the global metabolome in the probenecid exposure (furosemide coadministered with probenecid) arm.

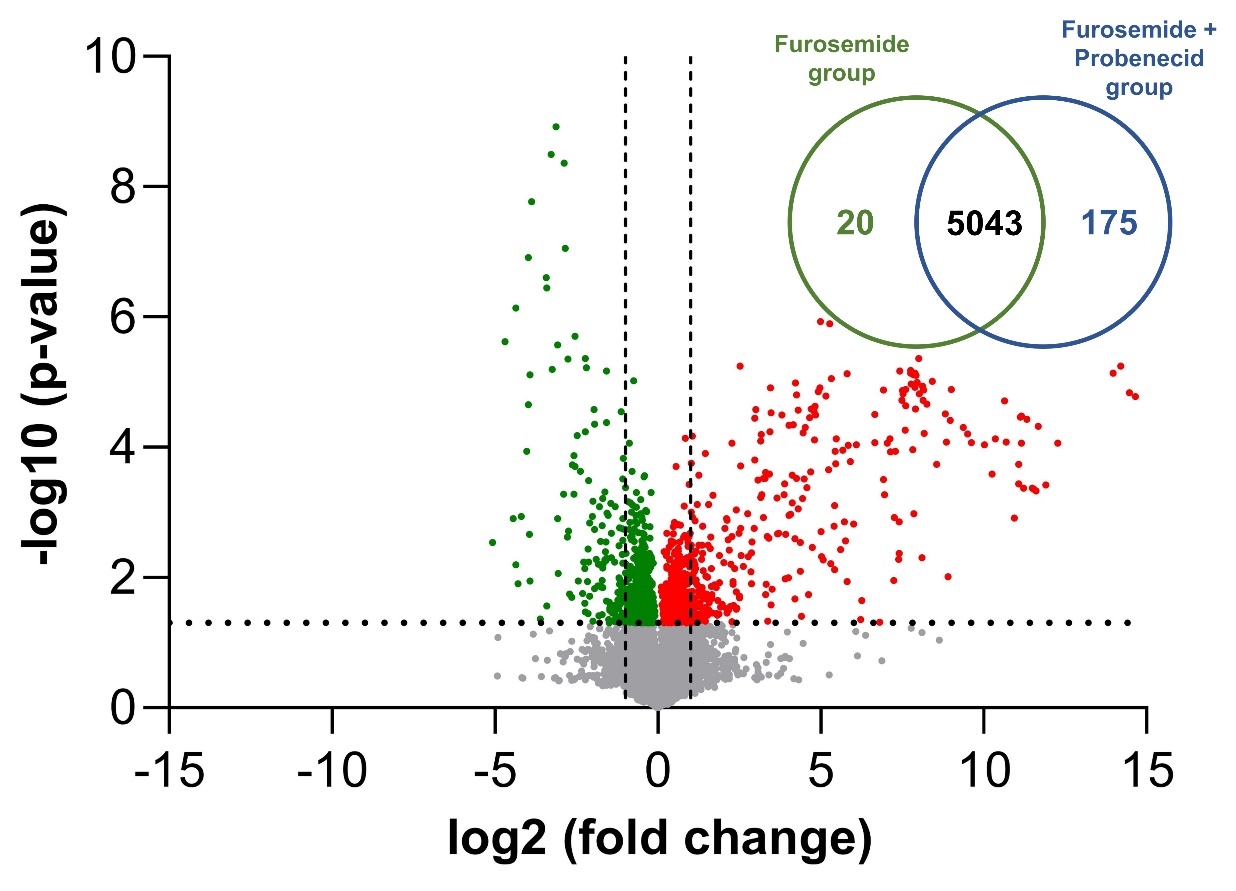

**Supplementary Figure 2. Comparison of *m/z* features detected in pooled plasma (1, 1.5, 2 h) in negative ionization mode in the baseline and probenecid exposure arms.** Venn diagram showing the total number of unique and common *m/z* features detected in the baseline and probenecid exposure arms. Volcano plot of the detected features, with dots representing *m/*z features that were either increased or decreased in pooled plasma in the presence of probenecid. Red and green dots indicate features that were significantly higher or lower (p-value < 0.05) in pooled plasma on probenecid exposure, respectively. The horizontal and vertical lines represent the p-value (< 0.05) and fold change (> or < 2-fold), respectively.

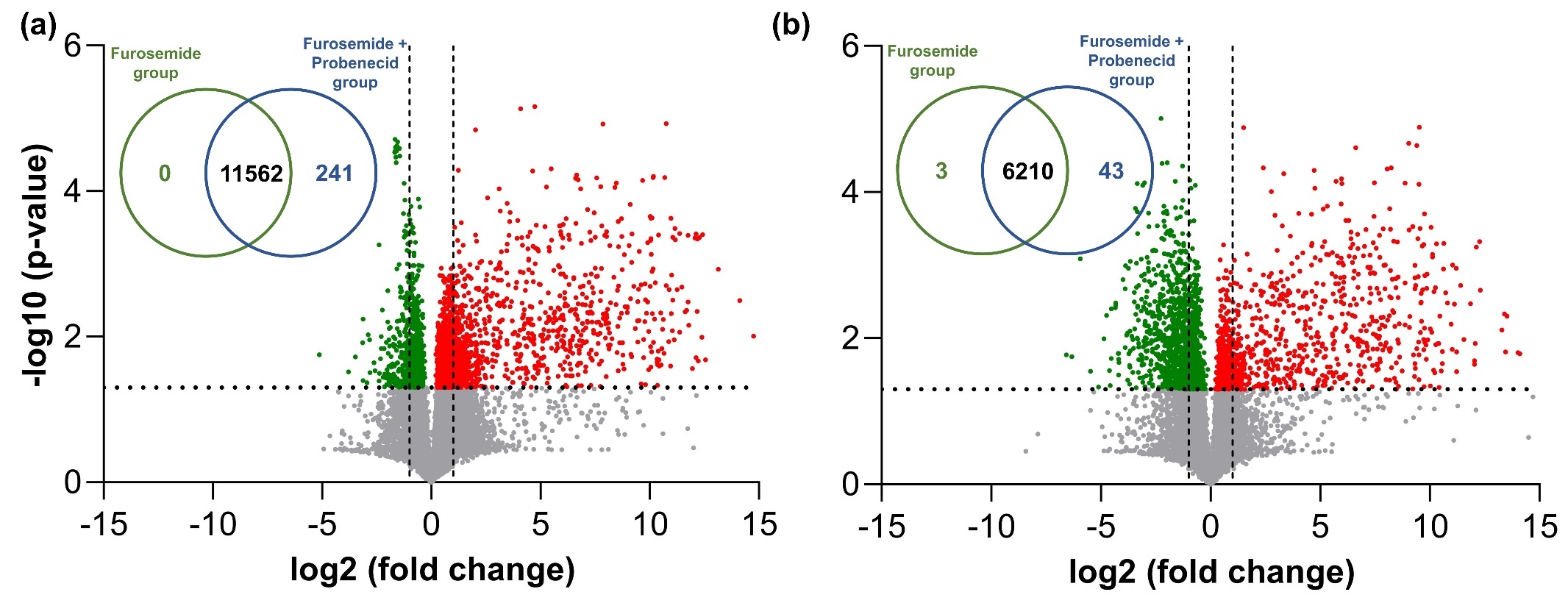

**Supplementary Figure 3. Comparison of *m/z* features detected in urine (0-4 h) in (a) positive and (b) negative ionization modes in the baseline and probenecid exposure arms.** Venn diagrams showing the total number of unique and common *m/z* features detected the in baseline and probenecid exposure arms in (a) positive, and (b) negative ionization modes. Volcano plots of the detected features, with dots representing *m/*z features that were either increased or decreased in urine in the presence of probenecid in (a) positive, and (b) negative ionization modes. Red and green dots indicate features that were significantly higher or lower (p-value < 0.05) in urine on probenecid exposure, respectively. The horizontal and vertical lines represent the p-value (< 0.05) and fold change (> or < 2-fold), respectively.

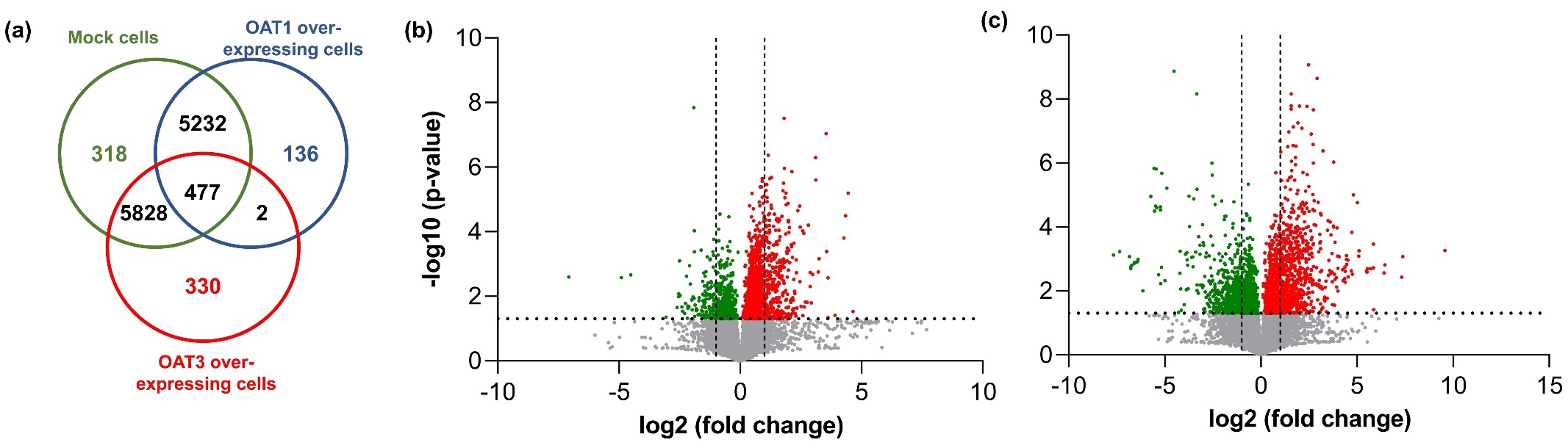

**Supplementary Figure 4.** **Comparison of *m/z* features concentrated in mock and OAT1 or OAT3 over-expressing cells following *in vitro* uptake assays in negative ionization mode.** (a) Venn diagram showing the total number of unique and common *m/z* features concentrated in mock cells, OAT1 over-expressing cells, and OAT3 over-expressing cells. Volcano plots of the concentrated features, with dots representing *m/*z features that showed either higher or lower uptake in (b) OAT1 over-expressing cells or (c) OAT3 over-expressing cells, compared to mock cells. Red and green dots indicate features with significantly higher or lower uptake (p-value < 0.05) in transporter over-expressing cells, respectively. The horizontal and vertical lines represent the p-value (< 0.05) and fold change (> or < 2-fold), respectively.

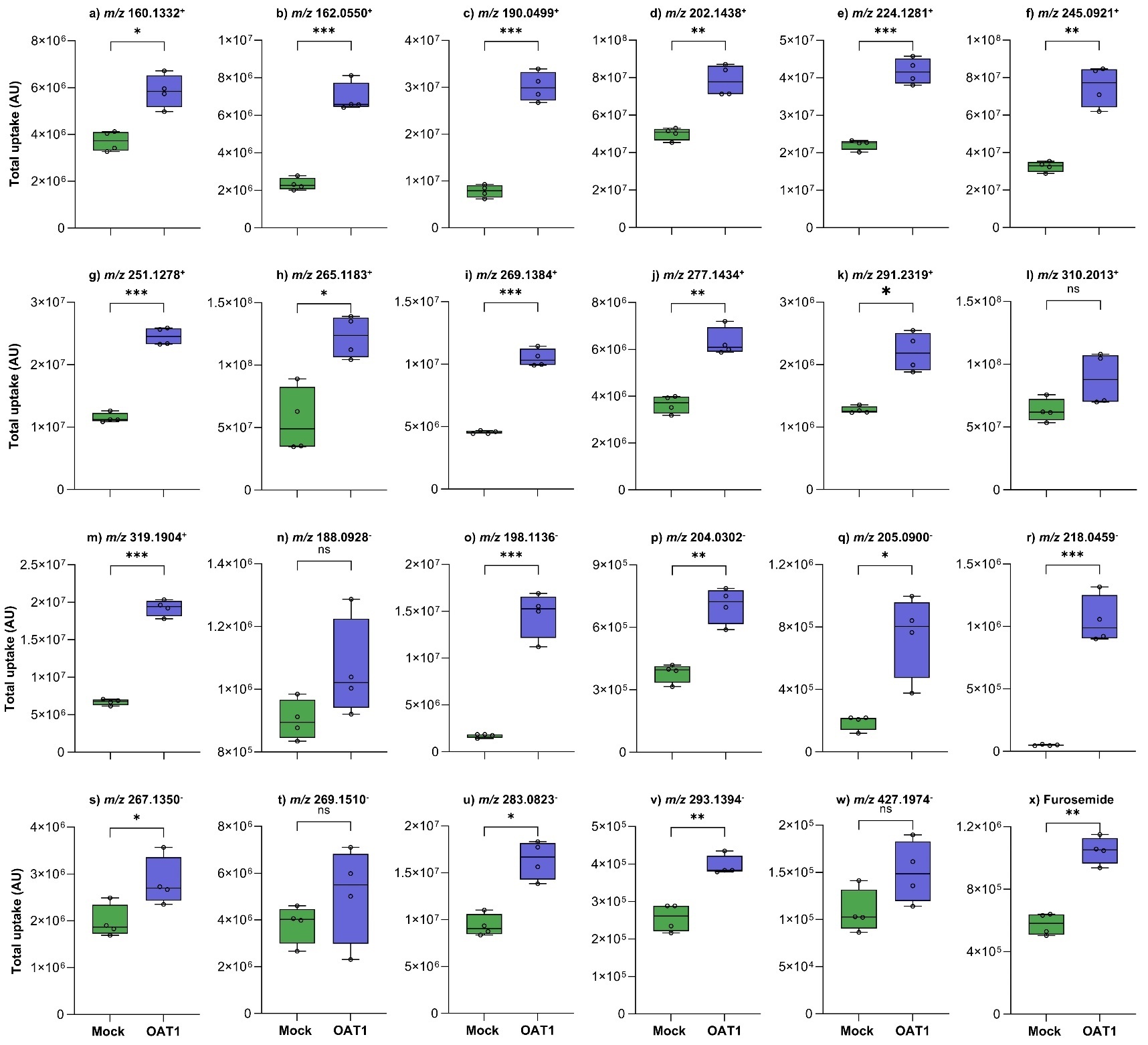

**Supplementary Figure 5.** **Uptake of validated OAT1/3 biomarkers and furosemide in OAT1 over-expressing cells compared to mock cells.** Box and whisker plots of validated OAT1/3 biomarkers (a-w) and furosemide (x) with the y-axis representing total uptake (peak intensity) of the *m/z* feature in mock (green box) and OAT1 over-expressing (blue box) cells. Whiskers in the box-and-whisker plots indicate the range, and the top, middle and bottom lines of the boxes represent the third quartile, median, and first quartile values, respectively. Open circles represent replicates (n=4). Total uptake (peak intensity) in mock cells and OAT1 over-expressing cells were compared using the unpaired t-test. p-value < 0.05 (*), < 0.001 (**), and < 0.0001 (***). AU = Arbitrary Units; OAT1 = Organic anion transporter 1; ns = non-significant; *m/z*: Mass-to-charge ratio with positive and negative sign representing the ionization mode.

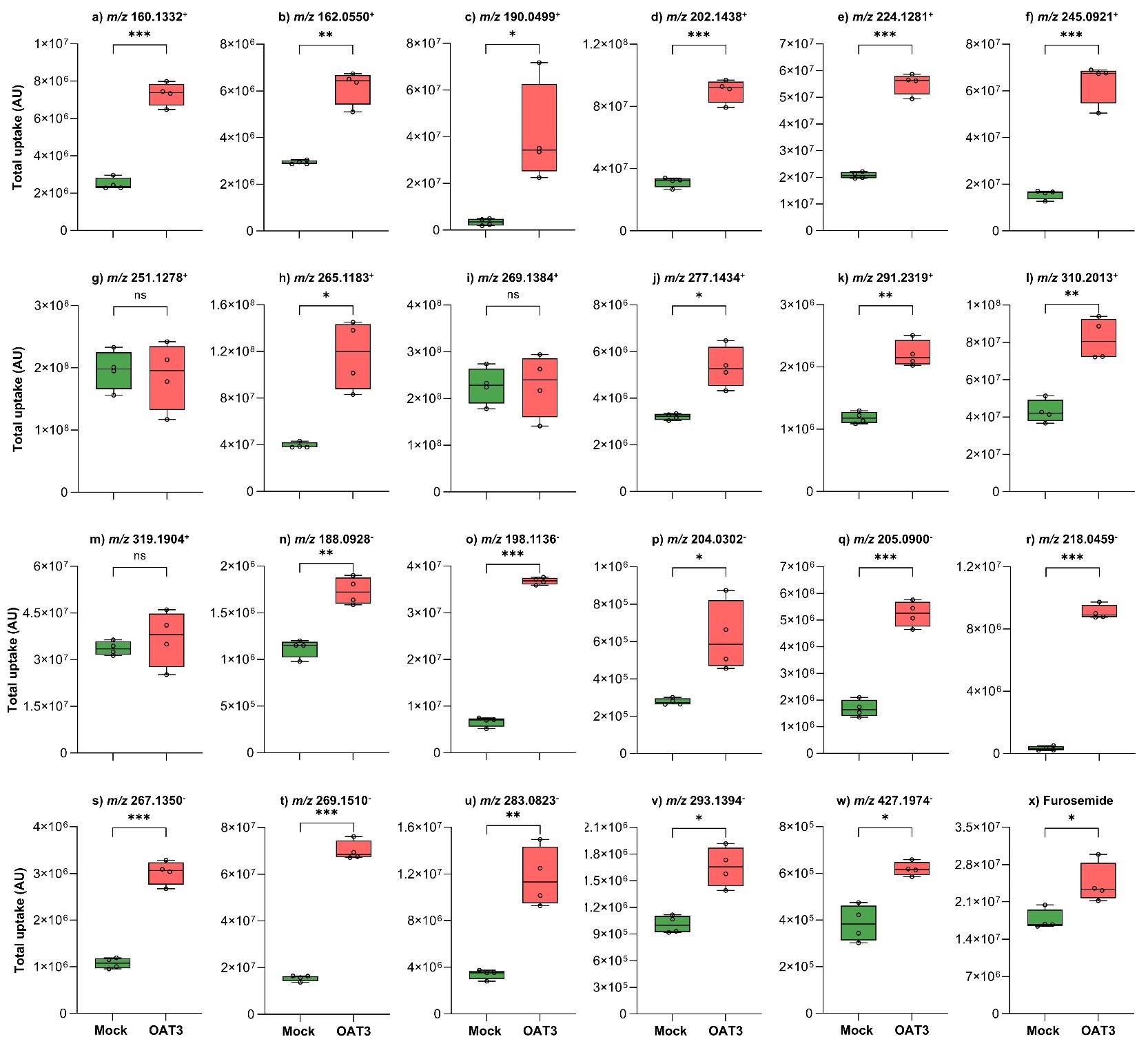

**Supplementary Figure 6. Uptake of validated OAT1/3 biomarkers and furosemide in OAT3 over-expressing cells compared to mock cells.** Box and whisker plots of validated OAT1/3 biomarkers (a-w) and furosemide (x) with the y-axis representing total uptake (peak intensity) of the *m/z* features in mock (green box) and OAT3 over-expressing (red box) cells. Whiskers in the box-and-whisker plots indicate the range, and the top, middle and bottom lines of the boxes represent the third quartile, median, and first quartile values, respectively. Open circles represent replicates (n=4). Total uptake (peak intensity) in mock cells and OAT3 over-expressing cells were compared using the unpaired t-test. p-value < 0.05 (*), < 0.001 (**), and < 0.0001 (***). AU = Arbitrary Units; OAT3 = Organic anion transporter 3; ns = non-significant; *m/z*: Mass-to-charge ratio with positive and negative sign representing the ionization mode.

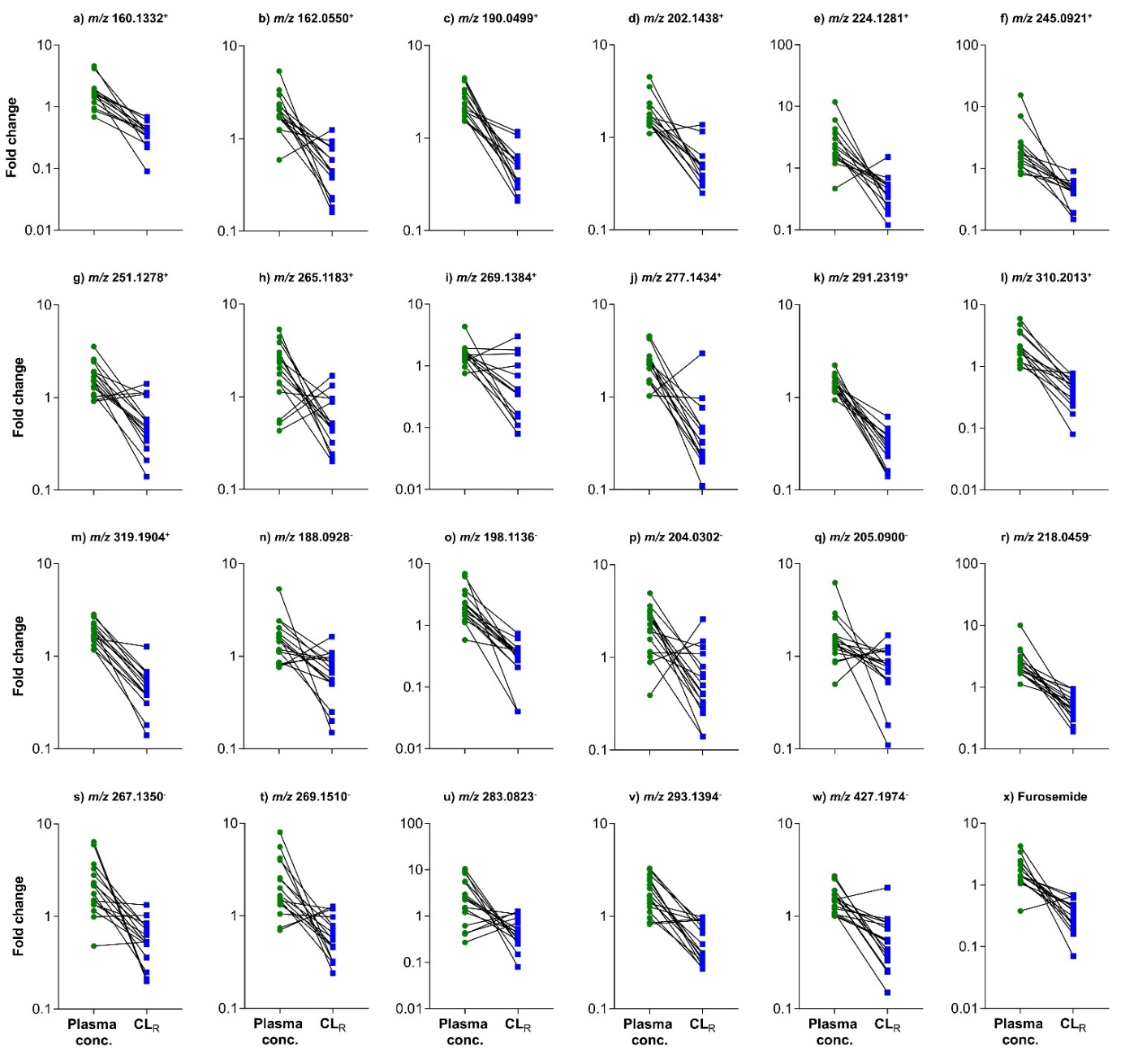

**Supplementary Figure 7.** **Fold change in pooled plasma concentration and apparent renal clearance (CL_R_) of validated OAT1/3 biomarkers and furosemide in the presence of probenecid.** Line and symbol plots of validated OAT1/3 biomarkers (a-w) and furosemide (x) with fold change on y-axis representing the ratio of pooled plasma concentration (green circles) or CL_R_ (blue squares) of the features in probenecid exposure arm versus baseline arm in healthy adult participants (n=16). Ratio = Pooled plasma concentration or CL_R_ in the probenecid exposure arm / Pooled plasma concentration or CL_R_ in the baseline arm. *m/z*: Mass-to-charge ratio with positive and negative sign in superscript representing the ionization mode.

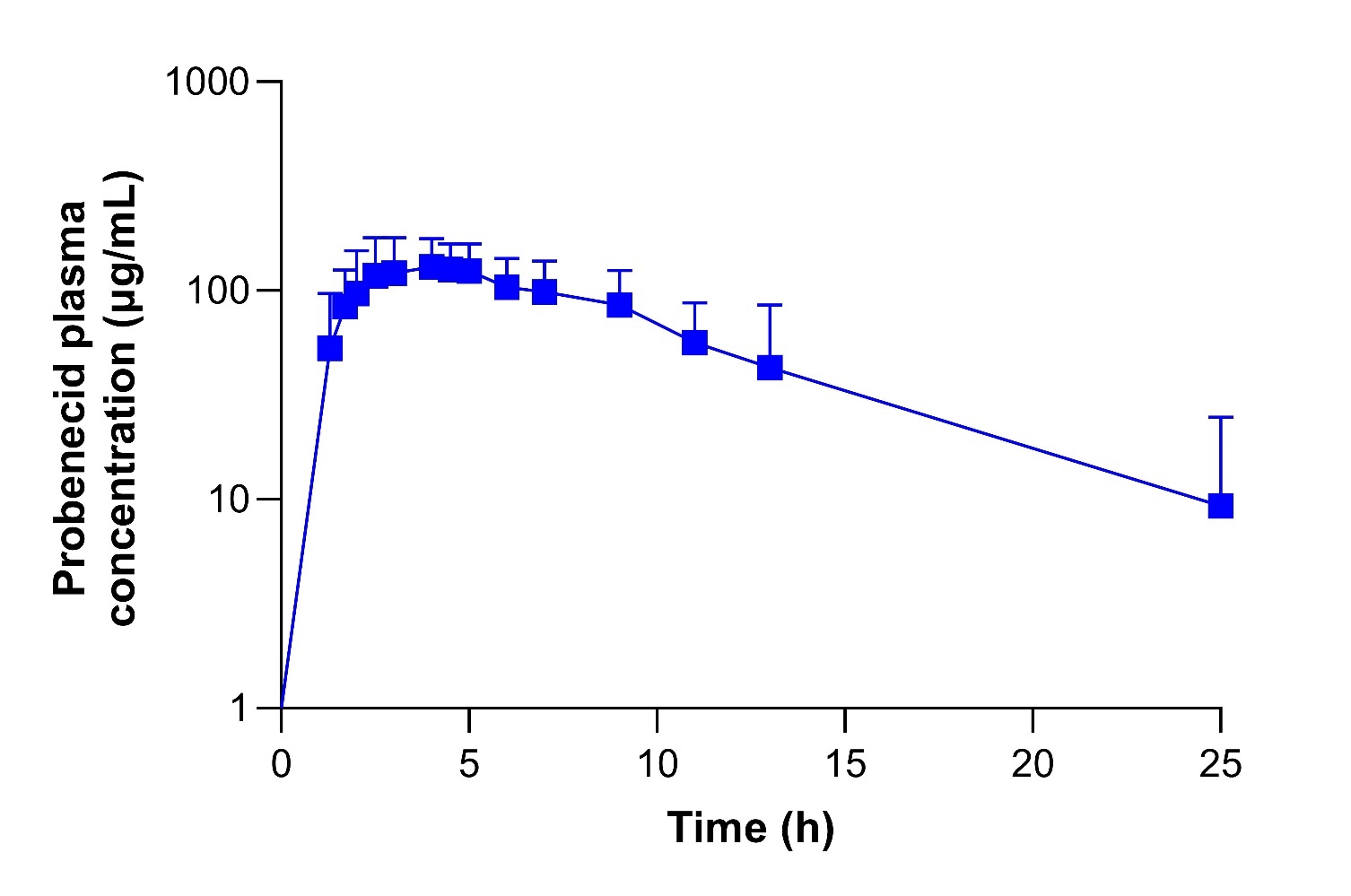

**Supplementary Figure 8.** **Pharmacokinetic profile of probenecid.** Average plasma concentration *versus* time profile of probenecid in healthy adult participants (n=16) in the probenecid exposure (blue squares, F+P) arm. The average data represent geometric means, with error bars indicating standard deviation. F+P = furosemide coadministered with probenecid (probenecid exposure arm).

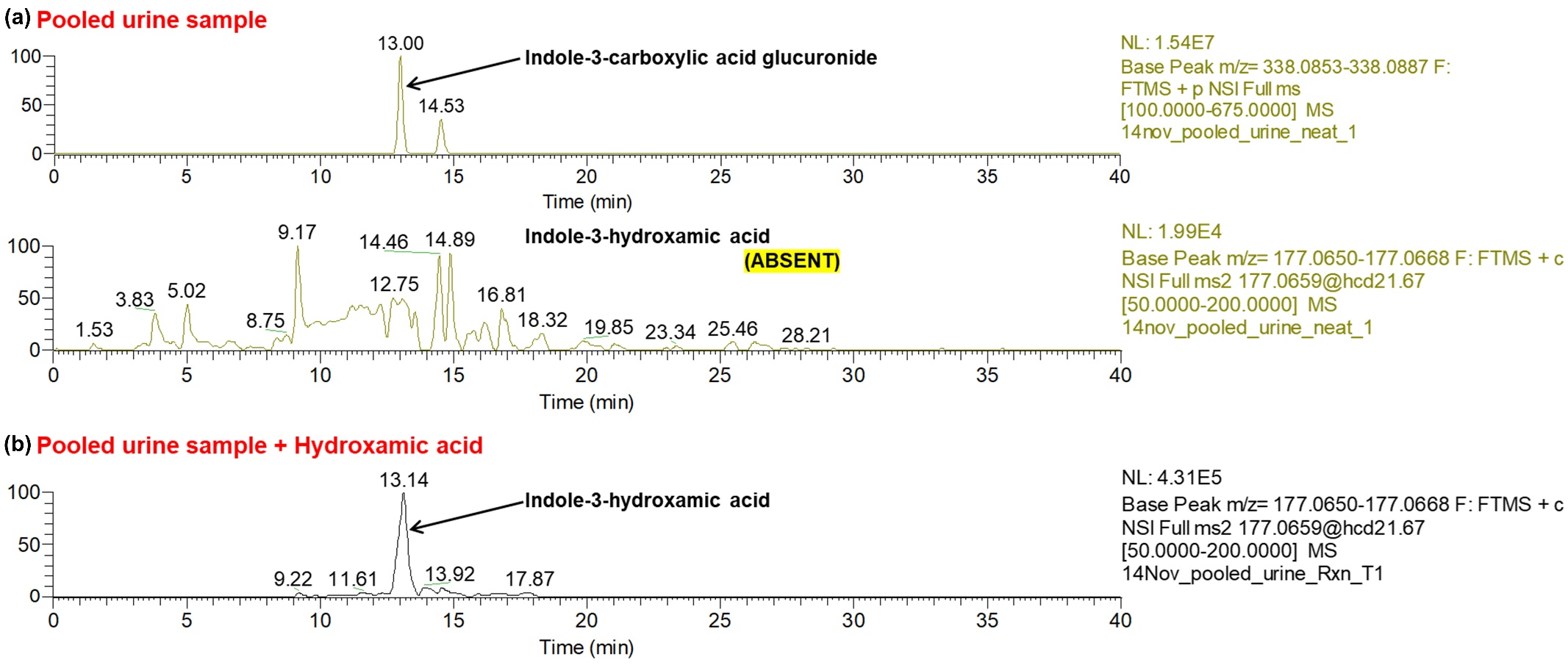

**Supplementary Figure 9. Derivatization of indole-3-carboxylic acid glucuronide (I3CAG) to indole-3-hydroxamic acid.** (a) The presence (top) and absence (bottom) of I3CAG and indole-3-hydroxamic acid in the pooled urine sample. (b) The formation of indole-3-hydroxamic acid from I3CAG on incubating the pooled urine sample with 50% aqueous hydroxylamine solution. Indole-3-hydroxamic acid was confirmed by MS/MS fragmentation-based structure characterization (data not shown).

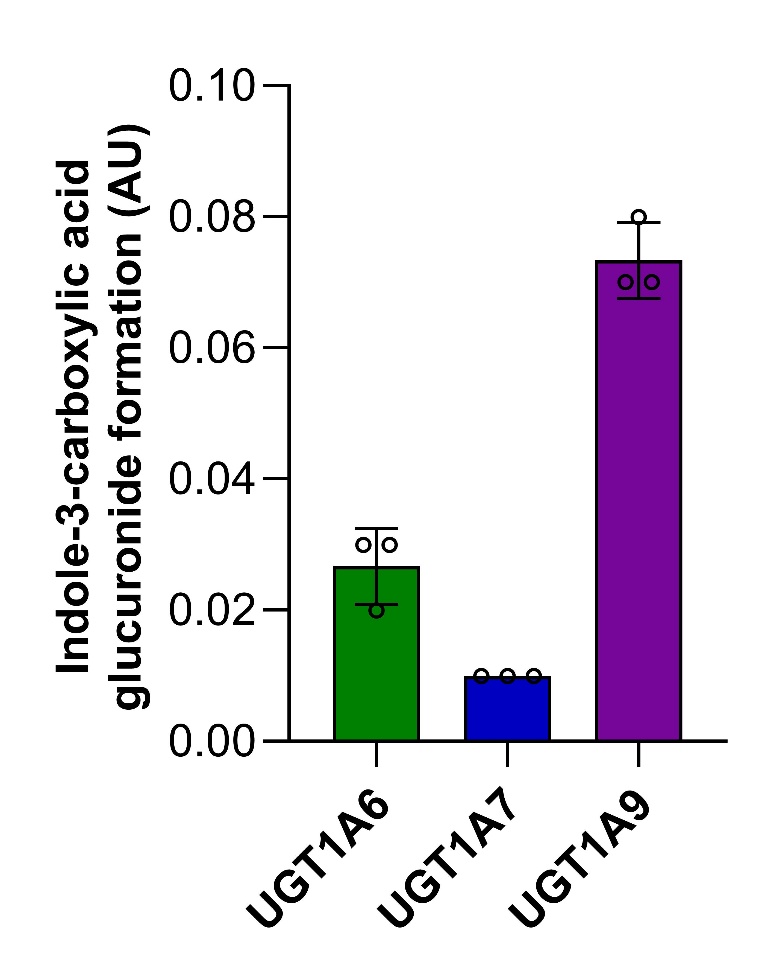

**Supplementary Figure 10.** **UGT isoforms involved in the formation of indole-3-carboxylic acid glucuronide (I3CAG).** Glucuronidation of I3CA by recombinant UGT1A6, UGT1A7 and UGT1A9 (n=3). A total of 13 recombinant UGT enzymes were tested for their ability to metabolize I3CA to its glucuronide. I3CAG formed by UGT1A1, UGT1A3, UGT1A4, UGT1A8, UGT1A10, UGT2B4, UGT2B7, UGT2B10, UGT2B15, and UGT2B17 was below the lower limit of quantification and is therefore not shown here. AU: Arbitrary Units (calculated by dividing the I3CAG peak area with testosterone glucuronide-d3 peak area).

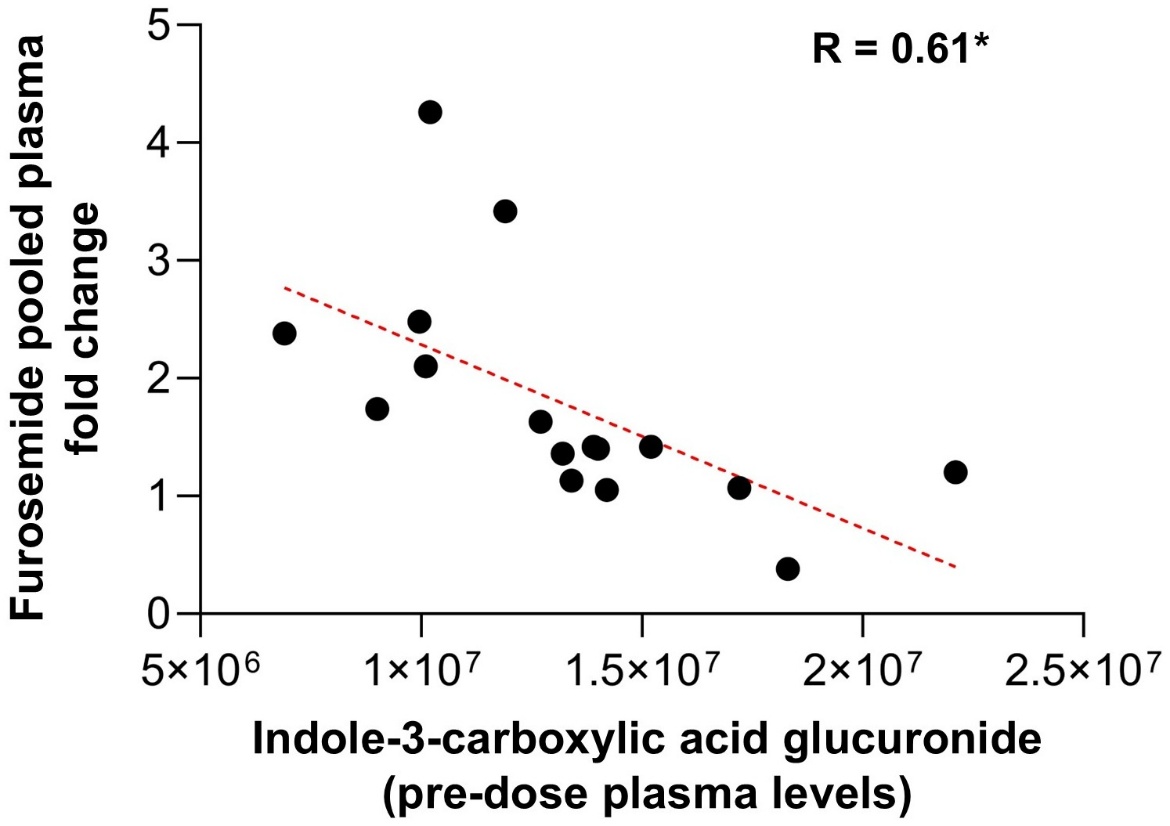

**Supplementary Figure 11.** **Correlation of indole-3-carboxylic acid pre-dose (0 min) plasma levels with furosemide pooled plasma fold change in healthy adult participants (n=16).** p-value < 0.05 (*).

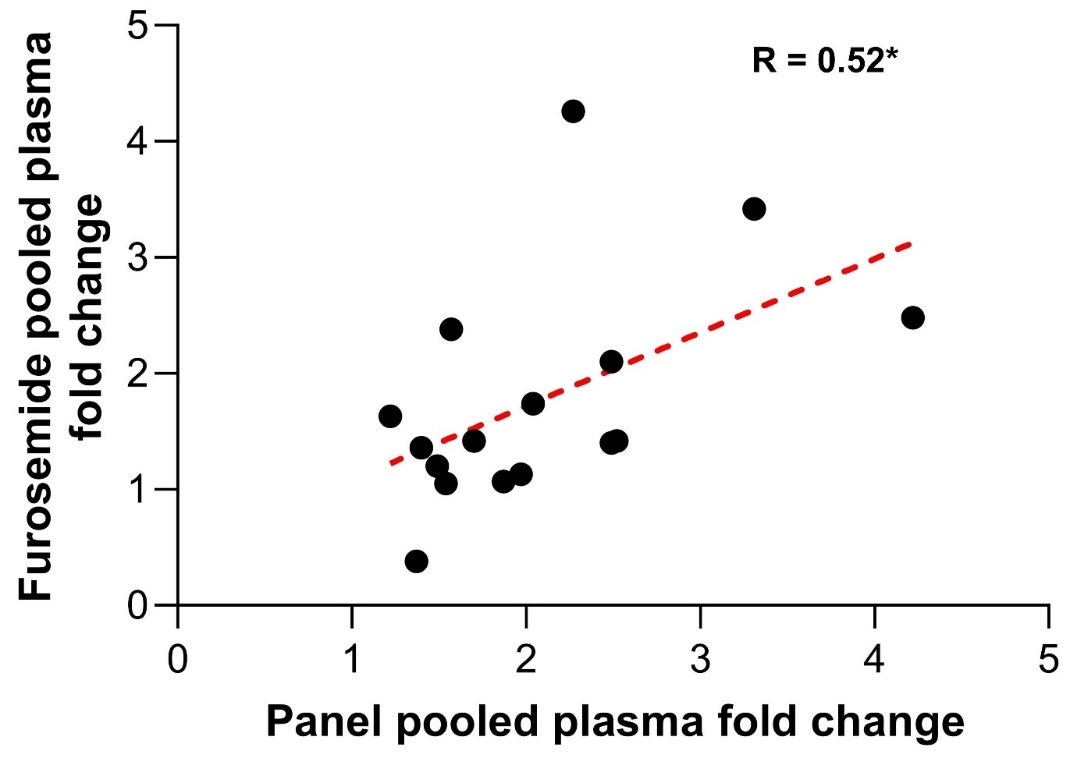

**Supplementary Figure 12. Utility of a panel of top six validated OAT1/3 biomarkers in predicting interindividual variability in probenecid-mediated changes in furosemide plasma concentrations.** Correlation between the average pooled plasma fold change of a panel of top six OAT1/3 biomarkers and furosemide pooled plasma fold change in healthy adult participants (n=16). p-value < 0.05 (*).
